# Flow Orchestrated Regulatory Genomics Engine (FORGE): A Configurable Nextflow Pipeline for End-to-End snMultiome Analysis

**DOI:** 10.64898/2026.09.10.750690

**Authors:** Luis E Solano, Negin Rahimzadeh, Zechuan Shi, Vivek Swarup

## Abstract

Single-nucleus multiome assays jointly profile gene expression and chromatin accessibility, yet their analysis typically requires bespoke chaining of modality-specific tools, creating barriers to reproducibility, scalability, and regulatory interpretation. We present FORGE (Flow Orchestrated Regulatory Genomics Engine), a configurable workflow that automates standalone snRNA-seq and snATAC-seq analysis, integrates the pair through complementary linear and nonlinear latent-variable models, and carries them through regulatory-network inference and differential testing. We evaluated FORGE on four human and mouse datasets spanning blood, brain, and kidney and two multiome chemistries, including a twelve-sample CRND8 Alzheimer’s disease cohort. We report cross-modal agreement alongside missing-modality reconstruction and an accounting of computational cost. In the Alzheimer’s cohort, FORGE nominated a glial Mef2c-associated program defensible across expression, co-accessibility, footprinting, and eRegulon evidence. In human PBMC, FORGE’s layered evidence models also provide nuanced interpretations that largely corroborate previously published regulatory links while also proposing an additional CD83 myeloid module.

**MOTIVATION:** Single-nucleus resolution multiome (snMultiome) assays concurrently profile gene expression and chromatin accessibility in the same nucleus. Yet regulatory inference from such analyses are difficult to scale, audit, and reproduce; moreover, as a field, snMultiomics and its’ toolset remains far from standardized. To address these challenges we developed FORGE, a configureable Nextflow workflow that carries paired data from raw counts and fragments through regulatory network inference with a comprehensive differential testing suite. Execution is containerized, tracks provenance, robust to interruption, optimized for cluster-based compute environments, and allows for nuanced customization of specific processes.

## INTRODUCTION

Single-cell and single-nucleus assays increasingly enable joint measurement of transcriptional and chromatin states, creating opportunities to connect cell-type-specific gene expression with regulatory architecture. In practice, however, multiome analysis commonly requires bespoke combinations of Bash, R, and Python scripts, with substantial variation in software versions, intermediate representations, quality-control decisions, and provenance. Workflow managers such as Nextflow (1) and Snakemake (2) can improve reproducibility, but existing community workflows are substantially more mature for snRNA-seq than for snATAC-seq or paired RNA-ATAC snMultiome analysis. For snRNA-Seq, epigen (3), bollito (4), and nef-core/scrnaseq (5) provide scalable automated processing (Table 1). For snATAC-Seq, only scATAC_snakemake (6) and scATAC-pipe (6) offer comparable architecture (Table 1). MAESTRO (7) stands as the sole dated resource addressing both modalities, however, it performs no analyses downstream of annotation and integration; most critically it lacks machinery for convergent regulatory inference (Table 1) which renders the most powerful component of snMultiome moot. Consequently, while these available resources address piecemeal portions of the problem, no peer-reviewed automated pipeline currently provides a reproducible end-to-end architecture extending from independent modality quality control (QC) through joint modeling, differential analysis, and regulatory inference.

**Table 1.** Comparison of automated workflow architecture capabilities supporting snMultiome analysis.

| Capability | FORGE<br>(this work) | MAESTRO | scGLUE | snRNA-seq<br>workflows<sup>a</sup> | snATAC-seq<br>workflows<sup>b</sup> |
| --- | --- | --- | --- | --- | --- |
| **Data modalities and integration** |  |  |  |  |  |
| Paired snMultiome (RNA + ATAC) | Y | Y | Y | N | N |
| snRNA-seq analysis (QC, integration, DGE) | Y | Y | Y | Y | N |
| snATAC-seq analysis (QC, peaks, annotation) | Y | Y | Y | N | Y |
| Linear integration (MOFA+) | Y | N | N | N | N |
| Non-linear integration (MultiVI) | Y | N | P | N | N |
| Cross-modal imputation / reconstruction | Y | N | Y | N | N |
| **Differential and regulatory analysis** |  |  |  |  |  |
| Differential RNA | Y | N | Y | Y | N |
| Differential ATAC | Y | N | P | N | Y |
| Co-accessibility (CCAN;<br>Cicero) | Y | N | N | N | Y |
| TF activity and footprinting<br>(chromVAR, scPrinter,<br>MSFP) | Y | N | P | N | P |
| Enhancer rewiring | Y | N | N | N | P |
| DORCs (correlated peak-gene links) | Y | N | P | N | N |
| eRegulons (SCENIC+) | Y | N | P | N | N |
| Gene regulatory network<br>inference | Y | N | Y | P | N |
| **Workflow engineering and reproducibility** |  |  |  |  |  |
| End-to-end reports and<br>visualizations | Y | Y | Y | P | Y |
| Workflow orchestration | Y | Y | N | Y | Y |
| Containerized execution | Y | Y | N | Y | Y |
| Checkpointing and<br>resumption | Y | Y | P | Y | Y |
| Provenance and audit trail | Y | Y | N | Y | Y |
| HPC / cloud scalability | Y | Y | Y | Y | Y |

Comparisons reflect features described in public documentation. Y, full support; P, partial support; N, not available. DGE, differential gene expression; DORC, domain of regulatory chromatin; HPC, high-performance computing; MSFP, multi-scale footprinting; snATAC-seq, single-nucleus assay for transposase-accessible chromatin sequencing; snRNA-seq, single-nucleus RNA sequencing; TF, transcription factor.
<sup>a</sup>Column reflects the union of capabilities across epigen, bollito, and the nf-core/scrnaseq workflow; no single workflow in this column provides all capabilities marked Y.
<sup>b</sup>Column reflects the union of capabilities across scATAC\_snakemake and the scATAC-pipe Nextflow workflow; no single workflow in this column provides all capabilities marked Y.

Here we present the Flow Orchestrated Regulatory Genomics Engine (FORGE), a configurable Nextflow architecture for standalone snRNA-seq, standalone snATAC-seq, and paired RNA-ATAC snMultiome analysis in human and mouse datasets (Table 1). FORGE does not replace the statistical methods it orchestrates; rather, its methodological contribution is to connect established tools within a versioned, containerized, resumable workflow with explicit provenance and consistent handoffs between transcriptomic, epigenomic, integration, and regulatory-analysis layers. We evaluate FORGE across four datasets spanning two species, three tissues, two snMultiome chemistries, and both single-condition and multisample comparative experimental designs. A central design principle is evidence convergence: candidate regulatory relationships can be evaluated across independent layers including expression, accessibility, motif activity, footprinting, co-accessibility, domains of regulatory chromatin (DORC), and eRegulons rather than being inferred from a single computational output.

FORGE’s package selection follows two constraints: every hand-off between stages is an AnnData/.h5ad object, and the RNA and ATAC arms must be able to run independently of one another. On the RNA side, scvi-tools supplies scVI, scANVI and MultiVI under one API, so batch correction, reference-guided label transfer, and the joint RNA+ATAC embedding share a single count-based probabilistic framework(8–10). Importantly, MultiVI imputes a missing modality (9), a function we demonstrate compatible with FORGE architecture by reproducing the publication’s held-out masking sweep. MOFA+ provides orthogonal convergent evidence through linear factor analysis, offering interpretable latent factors that reveal coordinated variation across RNA and ATAC (11) without the non-linearity of deep learning methods; its sparse factor loadings identify which genes and peaks drive each factor, and its factor space is used independently to assess embedding cohesion (Layer L2). CellBender precedes these tools because ambient RNA is enriched for the same highly expressed genes that define cell identity; its’ generative remove-background model returns posterior-corrected counts that remain integer-valued and therefore valid input to the count likelihoods downstream (12). For those unable to leverage the higher GPU-compute demands of reference-guided label transfer, CellTypist replaces the poorly reproducible stage of manual marker-gazing, with versioned logistic-regression models baked into the image (13). scanpy facilitates the steps between QC and clustering (14) while structuring the AnnData model that makes the persisting .h5ad currency possible. On the chromatin side, SnapATAC2 carries fragments through QC, tile and peak matrices, differential accessibility, and coverage export (15) entirely in Python, thereby avoiding potentially messy formatting conversions from R-based library usage into downstream ingesters. MACS3 is called per cell-type group, and merged into fixed-width non-overlapping peaks as chromVAR-style deviation computation and cross-cell-type coordinate comparison require (16,17). scATAnno annotates chromatin directly against a reference atlas (18), keeping ATAC labels independent of RNA. Areas where DNA and proteins bind, such as promoter and candidate cis-regulatory element (cCRE) genomic windows are relevant multi-scale footprint (MSFP) targets. scPrinter computes MSFP wherein the continuous Tn5-bias-corrected insertion-depletion score is calculated across many window widths simultaneously (19) and this MSFP scoring is saved to a single printer object per dataset. The same printer object also supplies the necessary structure to support a GPU chromVAR implementation and identification of background-calibrated DORCs peak-gene links when provided with paired expression data (19). SCENIC+, with pycisTopic, pycistarget, pySCENIC and MALLET, infers eRegulons (20–25). An eRegulon is comprised of a transcription factor (TF) together with the enhancers it binds and the genes those enhancers reach. The eRegulon is also the only one of the three regulatory layers that maps both modalities into a single testable TF to region to gene model; the layers are kept deliberately separate, and disagreements between them are reported. The R image exists because several required methods have no mature Python equivalent and all consume a Seurat object; the pipeline’s currency stays h5ad and enters R only where the best-available method lives there. Seurat supplies the differential-expression driver and the reference-projection machinery used for the independent annotation benchmark. MAST models the zero-inflated, bimodal per-cell expression distribution as a hurdle GLM with cellular detection rate as a covariate (26); its unit is the cell. The edgeR library is also present to enable pseudobulking to sample level with TMM normalization, robust dispersion estimation and quasi-likelihood F-tests (27). Both pseudoreplicate and pseudobulk approaches are reported and discussed at length. Next, weighted gene co-expression network analysis (hdWGCNA) makes co-expression analysis viable on sparse data by building metacells within cell type and sample before running WGCNA, converting thousands of per-gene calls into a few coordinated programs comparable across cell types and conditions (28). CellChat adds an intercellular layer, chosen over simpler ligand-receptor scoring because CellChatDB models multi-subunit complexes and cofactors explicitly and support condition contrasts (29). enrichR provides interpretation against Gene Ontology (GO) and KEGG, run separately on up-and down-regulated sets (30,31). The schard library facilitates reading .h5ad directly from R over HDF5 with no Python dependency, keeping the R processes hermetic. Finally, Cicero, through monocle3 with GenomicRanges, rtracklayer, and Gviz, estimates co-accessibility and groups links into cis-co-accessibility networks (CCANs) (32–36). Briefly, CCANs represent sets of peaks that open and close together, serving as data-derived cCRE-promoter units. It supplies the cCREs that MSFP footprinting targets, and because it requires no paired RNA, it enables promoter and cCRE interrogation inside the independent ATAC arm.

## RESULTS

### FORGE is Architecture for Automated snMultiome (RNA+ATAC) Analysis

FORGE utilizes a Nextflow architecture that accepts paired snMultiome (RNA + ATAC) file types and outputs a comprehensive analysis with end-to-end provenance; snRNA and snATAC analyses can also run as standalone workflows (Figure 1). RNA standalone analysis is comprised of ambient RNA cleanup (37), quantile based filtering (38), a batch-aware integration (39), cell annotation (13,40), differential gene expression (DGE) (41), functional enrichment (42,43), hdWGCNA (28), and cell communication inference (44) (Figure 1). ATAC standalone analysis is comprised of an intial per fragment quality control (QC) (45), a secondary quantile-based per-cell QC (45), a critical accessibility metrics quantification (45), peak calling (16,45), peak clustering (45), reference-based cell annotation (46), cis-coaccessibiliity network (CCAN) generation (34), evaluation of transcription factor (TF) motif activity (19,47), enhancer rewiring (19,34), and multiscale footprinting (MSFP) (19) (Figure 1). If sufficient cells meet QC and independent modality upstream analyses thresholds, cross-modal integration (48), and a two-way (linear MOFA+ and non-linear MultiVI) factor analysis are executed (49,50) (Figure 1). Following cross modal integration (49–51), users can rescue dropout cells via tuneable imputation (39,50), calculate SCENIC+ eRegulons (52), calculate DORCs (19), and generate gene regulatory networks (GRN) (52) (Figure 1).

**Figure 1.**
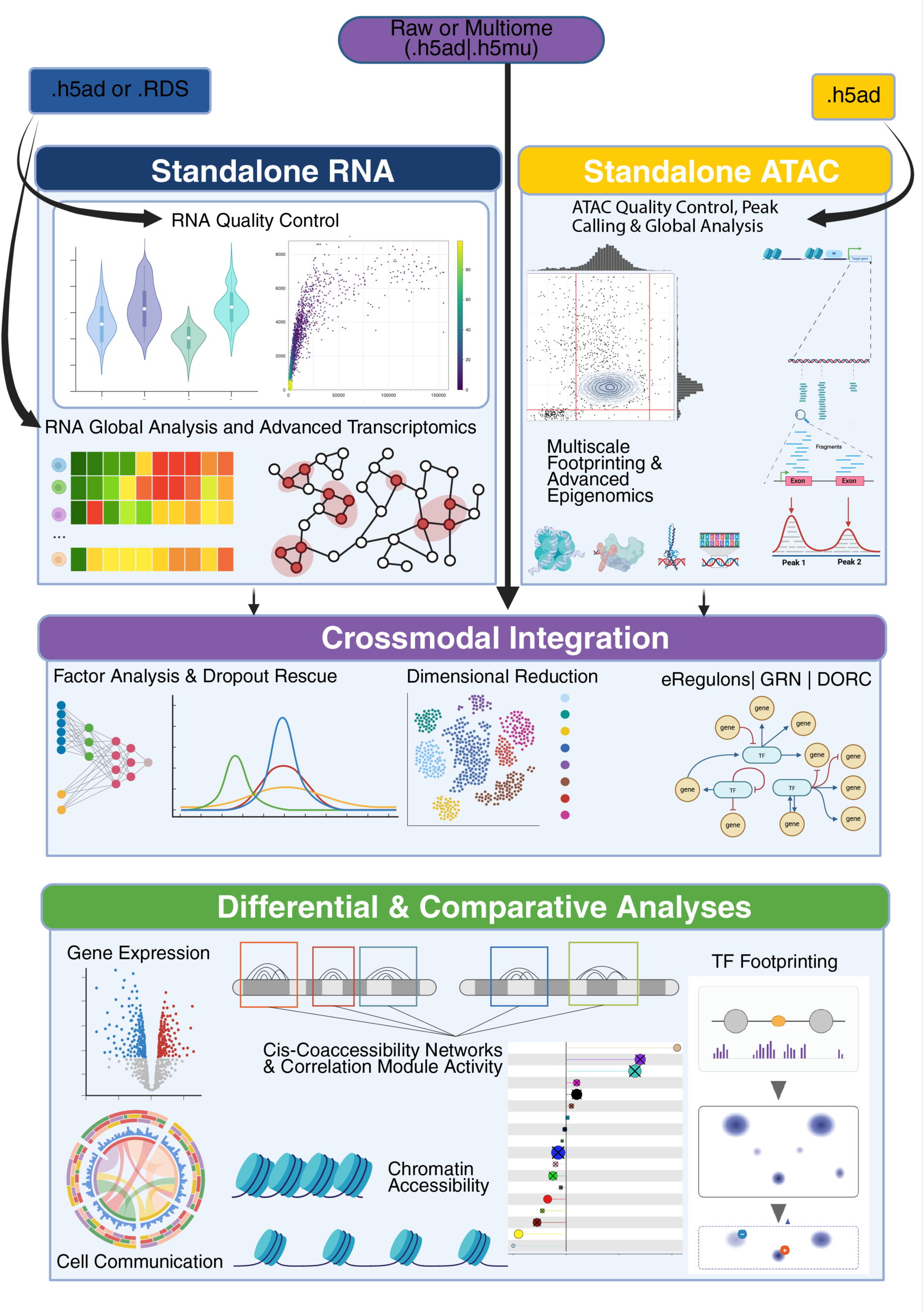
FORGE nextflow architecture automates scalable snMultiome analyses in mouse and human. Beginning from either standalone RNA or ATAC inputs, FORGE transparently and reproducibly standardizes and automates snMultiome analysis to output statistically robust results including modes for differential analyses. The standalone RNA mode performs QC, dimensional reduction, annotation, integration, and global statistical transcriptomic analyses. Notably QC steps are comprised of ambient transcript removal and quantile-based filtering. Global analyses include dimensional reduction and various subtype annotation refinement methods. The standalone ATAC mode performs QC, dimensional reduction, annotation, integration, and global statistical epigenomic analyses. When users provide paired modality data, the cross-modal integration modes are made available including factor analysis, dropout rescue, SCENIC+ eRegulon calcuation, gene regulatory network construction, and DORC computation. Specifically, both non-linear and linear factor analyses are available with non-linear embeddings offering cross-modal rescue of unpaired cells via imputation. With multiple samples and conditions, the differential modes enable statistically rigourous testing of gene expression, cell communication, chromatin accessibility, TF footprinting, and correlation module analyses.

For standalone RNA, users can enter the pipeline using .h5ad or .RDS files from pre-or post-ambient RNA removal, RNA QC, global analysis, or advanced transcriptomic analysis (Figure 1). While we recommend gpu-accellerated reference-based RNA cell annotations via SCANVI (39,40), users may elect for a CellTypist (13) model or a custom manually input marker based annotation. For standalone ATAC users, users can enter the pipeline from fragment files or post QC .h5ad files (Figure 1). Differential analysis modes for RNA include expression via MAST (41), cell communication via CellChat (44), and module activity via hdWGCNA (28) while differential analysis modes for ATAC include chromatin accessibility via snapATAC2 (45), peak calling via MACS3 (16,45), MSFP via scPrinter (19), and CCANs via Cicero (53) (Figure 1).

### FORGE Constructs Concordant Linear and Non-linear Paired Multiome Integrations Identifying Gene Expression Profiles with Orthogonal Chromatin Accessibility Evidence

10x Genomics profiled gene expression and chromatin accessibility from human peripheral blood mononuclear cell (PBMC) nuclei (54), providing a widely used benchmark for paired multiome analysis. FORGE retained 7,620 nuclei with paired RNA and ATAC measurements after modality-specific QC and generated independent annotations and joint representations for downstream evaluation (Figure 2A). We integrated the paired modalities using MOFA+ and MultiVI (49,50), which provide complementary linear and nonlinear latent representations with distinct modeling assumptions. Agreement between representations identifies biological structure that is robust to these distinct frameworks; conversely, discordant dimensions are treated as candidates for method-dependent or potentially nonlinear structure that require additional evaluation rather than as evidence of nonlinear biology by themselves (Figure 2B-D). Regulatory outputs were subsequently evaluated across independent evidence layers, including SCENIC+ eRegulons and scPrinter-derived DORCs (19,52) (Figure 2E-F). We additionally quantified computational resource requirements to distinguish the cost of core processing from optional high-cost regulatory analyses (Figure 2G; Table 2).

**Figure 2.**
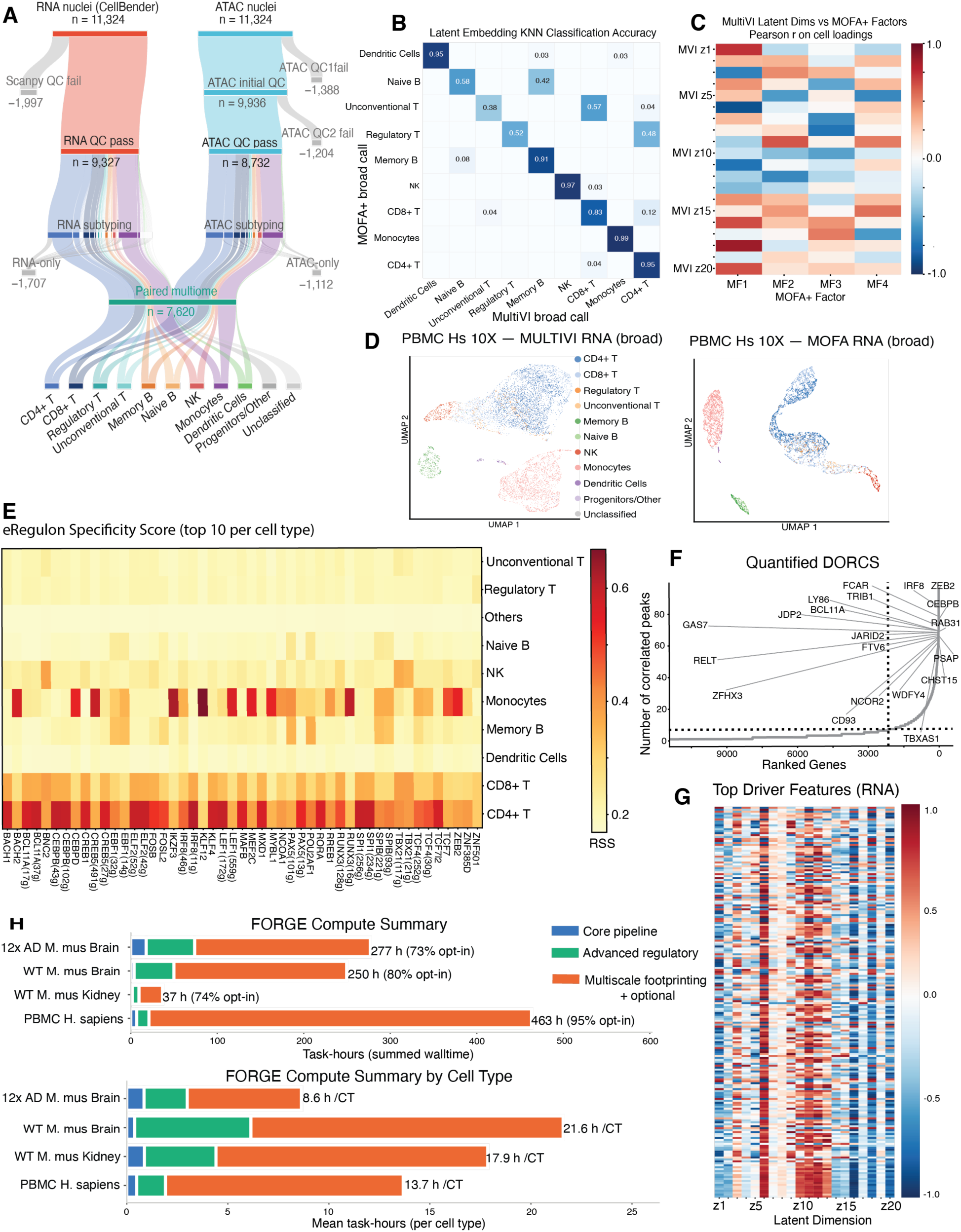
PBMC snMultiome inputs demonstrates identification of canonical cell types, associates factors derived from integrated embeddings with biological programs, and constructs gene regulatory networks justified via orthogonal computational methods. An example of a FORGE end-to-end analysis demonstrating QC outputs, integrated dimensional reduction, putative gene regulatory networks, and a quantitative summary benchmarking of computation. (A) Quantitative summary of FORGE QC steps which identify 7620 nuclei distributed among 11 broad cell types. (B) Minimal B cell and T cell subtyping errors dominate discrepancies in a KNN classification task to assess agreement between broad cell type annotations within orthogonal latent multiome embeddings. (C) Pearson correlations between MOFA+ factor cell loadings and MultiVI latent embedding dimension cell loadings show emergent agreement among orthogonal methods and illuminates opportunities for exploration where anti-correlations indicate non-linear biology. (D) UMAP visualizations of MultiVI and MOFA+ multiome latent embeddings colored by RNA annotation. (E) Various transcription factors plotted by their respective eRegulon’s rank bi-serial score show Monocytes and CD4+ T cells dominate the strongest PBMC signal for potential eRegulon-based mechanisms of action. (F) Domains of regulatory chromatin (DORC) orthogonally computed via scPrinter corroborate various eRegulon signals. (G) Gene expression signature associated with the top 20 MultiVI dimensions suggest varying levels of the sensitivity of specific MultiVI dimensions to changes in expression. (H) FORGE runtime quantitatively summarized by major processes and averaged over cell type. Each of the 4 validated datasests recorded compute time and resources consumed. The lower bar plot tracks compute costs per-cell-type to estimate how FORGE may scale for larger and more diverse datasets.

**Table 2.** Summary quantification of compute hours and memory resources.

| Process | Compute Hours | Peak Ram (GB) |
| --- | --- | --- |
| Enhancer footprinting per CT | 562.7 | 320 |
| Scprinter footprinting | 120.3 | 116 |
| Scprinter footprinting DIFF | 49.9 | 113 |
| MultiVI masking sweep | 40.4 | 163 |
| Enhancer footprinting per CT strip | 27.0 | 168 |
| Run Cellchat | 25.3 | 43 |
| Cicero full chromosome per CT | 19.4 | 5 |
| PyCisTopic prepare | 18.2 | 249 |
| Promoter MSFP per CT | 15.1 | 142 |
| HDWGCNA enrichment | 14.0 | 9 |
| Cellbender | 12.6 | 34 |
| scPrinter DORC | 12.5 | 149 |
| Rank enhancer strip genes | 10.9 | 1 |
| Composite enhancer viz | 9.2 | 2 |
| SCENICplus run | 8.9 | 376 |
| MultiVI gap fill | 6.9 | 629 |
| Build muData | <1.0 | 569 |
| MultiVI validate | <1.0 | 232 |
| Prepare Reference | <1.0 | 174 |
| Total | 953.8 | Max = 629 |

FORGE was developed and optimized on a high-performance computing server (Rocky 9.7 Blue Onyx). To report the required compute resources to run FORGE end-to-end, we quantified memory usage and total runtime via compute hours for each dataset globally (Figure 2H, nextflow.config files, and Table 2). Additionally, we also report per-average celltype runtime and compute hours to grant users an estimate of compute resources and time required to run FORGE at scale (Figure 2H, Table 1). We do note that the enhancer footprinting process’s default discovery mode utilizes architecture that fans out top TF candidates on uniquely called fine-grain cell types (Figure 2H, Table 2). Thus, enhancer footprinting can overwhelmingly dominate the runtime; deeply subtyped datasets or workflows where many TF’s are evaluated for enhancer footprinting, such as in the human PBMC example, will incur heavier compute burdens. When considered on an average compute-hours per celltype basis, the compute hours runtime is more comparable across datasets (Figure 2H, Table 2).

### FORGE Surfaces A Putative Mef2c Driven Program Demonstrating Native Multi-sample Scaling and Comparative snMultiomic Analyses

We applied FORGE to a public 10x Genomics multiome dataset of wild-type and CRND8 APP-overexpressing transgenic mouse brain (55), focusing on paired glial nuclei to assess convergence of independent regulatory evidence. Differential expression recovered established glial responses (56–58), while SCENIC+ identified a Mef2c-associated eRegulon with cell-type-specific activity differences. Complementary evidence from Mef2c expression, target accessibility, MultiVI factor z19, Cicero co-accessibility, chromVAR motif activity, and scPrinter footprinting supported this glial Mef2c program (Figure 3). SCENIC+ revealed activity shifts in astrocytes (LFC=1.22, p<0.05) and oligodendrocytes (LFC=1.91, p<0.05), with opposing shift in microglia (LFC=0.15, p<0.05) (Figure 3C). Mef2c expression was confirmed across glial types with >80% of canonical targets accessible (Figure 3C). MultiVI factor z19 explained TGvsWT variance from Mef2c regulation (rank biserial scores: microglia 0.14, astrocytes 0.19, oligodendrocytes 0.26), with 20 of 30 top RNA drivers as Mef2c targets with accessible motifs (Figure 3C). Cicero analysis identified Mef2c in driver-TF-gene-cCRE modules (Figure 3D) and TG/WT-associated co-expression modules (Figure 3E). ChromVAR z-scores revealed shared and distinct regulatory programs (Figure 3E). scPrinter footprinting on ENCODE/SCREEN-confirmed regions showed differential TF binding: increased Spi1 at Trem2’s microglia promoter, decreased Mef2c at Fras1’s astrocyte promoter, and increased Mef2c at oligodendrocyte Tmem151a promoter (Figure 3F); altered Mef2c binding at glial cCREs also evident (Figure 3F). Finally, Cellchat analysis revealed differential cell-cell communication across TGvsWT (Figure 3G).

**Figure 3.**
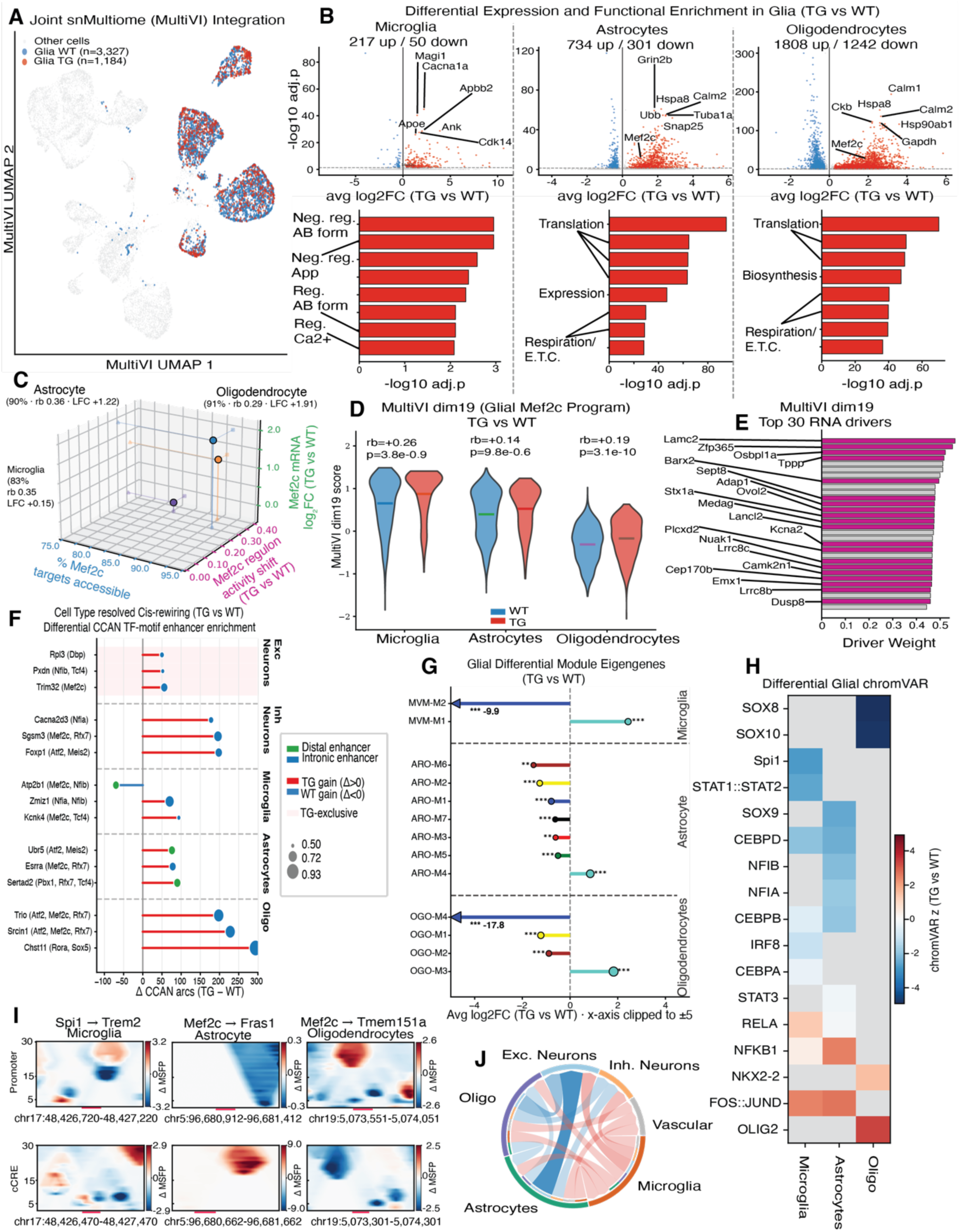
FORGE proposes a Mef2c driven regulatory program justified by cross-modal multiomic evidence in a transgenic AD mouse model. After subsetting cells to a glial population, FORGE identifies a Mef2c regulatory program via orthogonal compute methods spanning statistically significant changes in gene expression, accessibility, transcription factor (TF) binding, and cisCRE wiring. (A) The joint snMultiome MultiVI embedding demonstrates FORGE properly clusters different cell populations and highlights glial cells to demonstrate the embedding is well integrated. (B) Differential expression analysis performed in glial populations demonstrate canonical AD DEGs in addition to Mef2c; functional enrichment analyses also highlight canonical pathways. (C) Mef2c expression (z-axis in green) combined with the proportion of accessible Mef2c targets (x-axis in blue) in Microglial, Astrocyte, and Oligodendrocyte populations confirm that expression and accessibility are consistent with the differential Mef2c eRegulon activity shift identified by SCENIC+ (y-axis in purple). (D) The MultiVI dim19 loading defines the Mef2c regulatory program in Microglia and Astrocytes. (E) Twenty of the top 30 RNA drivers of the MultiVI dim19 loading contain the Mef2c binding motif. (F) Differential CCAN analyses identifies a near-global increase in peak-gene links reported and stratified by cell types, cCRE location, and proportion of arcs with the named TF-motif enrichment. (G) Module correlation analysis via HDWGCNA identifies a dominant negative shift in statistically significant glial module eigengenes (p<0.05) with the turquoise module acting as the exception. (H) Differential chromVAR in glia indicates TG aligns with canonical AD mouse model TF activity. Heatmap cells colored according to a z-score. (I) Multiscale footprint plots of 150bp genomic windows surrounding gene promoter and cCRE regions for a TF-gene pair in microglia, astrocyte, and oligodendrocytes. Red coloration indicates a calculated TG footprinting score with a positive ratio compared to WT at the indicated coordinates (genomic position (x-axis, bp) and footprint size (y-axis, bp)); blue indicates a negative ratio compared to WT. Positive ratios are interpreted as a gain or increase in protein-gene binding whereas negative ratios suggest a loss or decrease in binding. (J) Cellchat uncovers evidence for astrocyte to excitatory neuron signaling among a generalized trend towards outbound communications from glial populations towards neural.

### Replicate-aware pseudobulk analysis silences the Mef2C regulatory signal, but reproduces constituent effect sizes

Importantly, the differential pseudoreplicate analyses just described identify putative regulatory relationships and nominate testable hypotheses; cell-level pseudoreplicate differential tests treat cells from the same animal as independent replicates and therefore risk inflating type 1 errors. We re-tested the TG-versus-WT contrast on pseudobulk libraries formed by summing raw counts within each (cell type, mouse), modelling age as a continuous covariate across the 12 animals (Figure 4). The re-analysis is essentially null with 340 of 219,519 gene-level and 76 of 943,015 peak-level tests reaching FDR < 0.05 (Figure 4A, 4B), reducing 17,804 MAST calls to 340. The collapse is not an implementation failure: an internal positive control run through identical machinery, contrasting microglial/immune against OPC/oligodendrocyte accessibility paired within mouse, recovers 33,759 of 65,079 peaks (51.9%, BCV 0.133; Figure 4A). Nor is it confined to per-gene testing. Rotation-based gene-set testing (fry) finds no direct activating eRegulon differentially active at FDR < 0.05 in any glial population, Mef2c included (p = 0.61 astrocytes, 0.64 oligodendrocytes, 0.19 microglia), while recovering 44 of 47 regulons on the control contrast (minimum p = 7.4e-12; Figure 4C); the competitive alternative camera, which corrects for the inter-gene correlation measured in these data (rho = 0.020-0.036), agrees (p = 0.23-0.44). With six animals per genotype distributed across three ages, this cohort is underpowered to evaluate condition effects at either the gene or program level. What the pseudobulk analysis does provide is an independent estimate of effect size, computed under a model that makes no independence assumption about cells. Across every gene MAST called, cell-level and replicate-aware log2 fold changes are concordant (pooled Pearson r = 0.61, 73.0% agreeing on direction, n = 17,733) (Figure 4D). Concordance is strongly cell-type dependent: it is high in glia (per-cell-type r = 0.82-0.95, 90% directional agreement) and close to chance in deep-layer excitatory neurons (r = 0.44, 65% directional agreement) (Figure 4D). The ATAC arm behaves the same way (r = 0.60, 95% directional agreement, n = 112; Figure 4E). Critically, the agreement holds on the gene set the regulatory model depends on: across the 2,506 genes of the direct activating Mef2c eRegulon, r = 0.92 in astrocytes (n = 316 MAST-called targets), 0.85 in oligodendrocytes (n = 876) and 0.93 in microglia (n = 91), with aggregate 88-98% directional agreement (Figure 4F). Effect sizes are systematically attenuated (regression slope 0.27; median absolute log2FC 0.76 falling to 0.23), consistent with the expected overstated magnitude inherent to cell-level tests; however, their sign and relative ordering are preserved.

**Figure 4.**
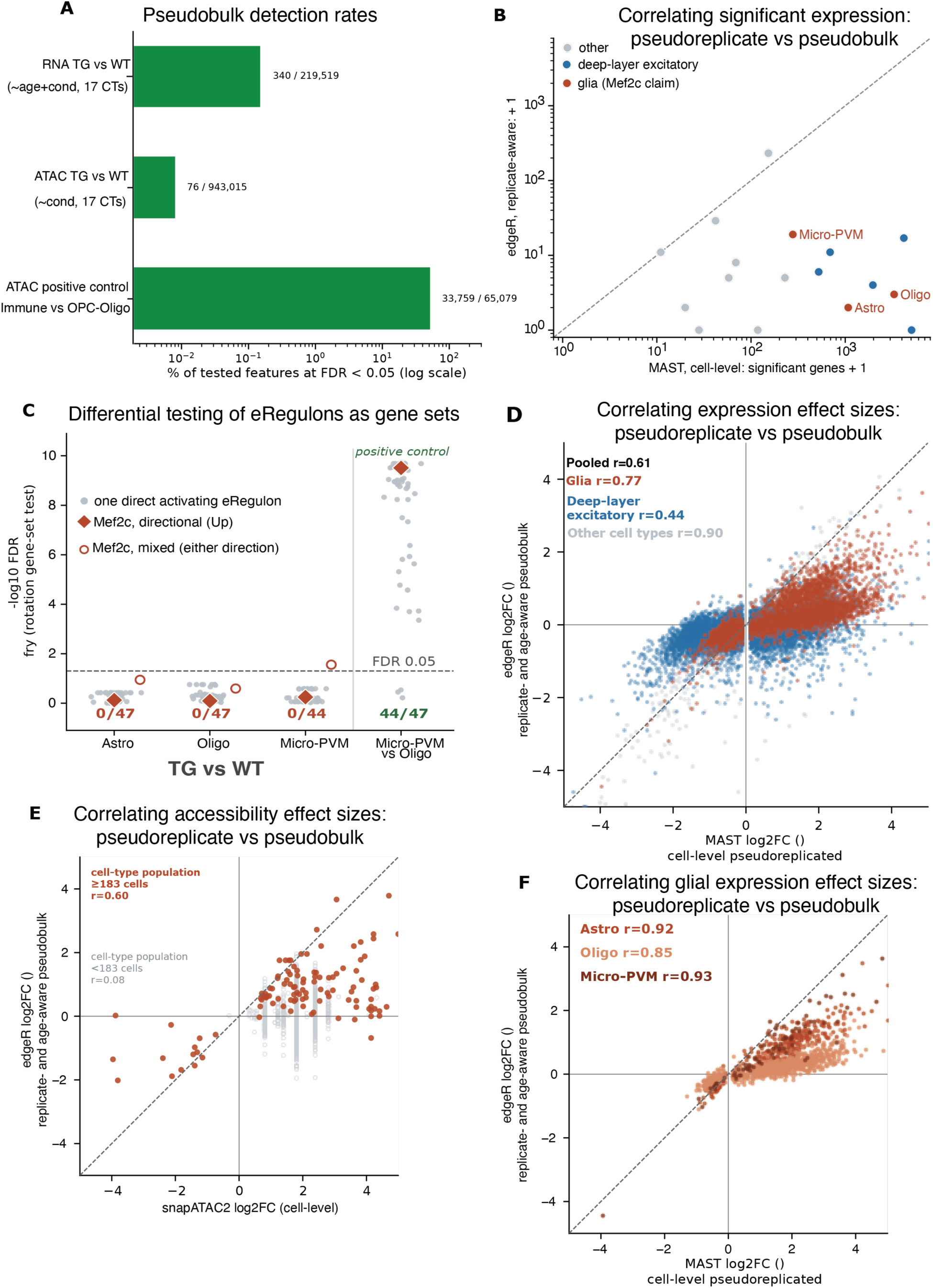
DEGs and DARs supporting Mef2c program collapse under replicate- and age-aware pseudobulk testing but preserve effect sizes and their rankings. A replicate- and age-aware pseudobulk analysis silences the single-cell pseudoreplicate condition signal at the gene and program levels; correlations surface agreement on effect size, offering orthogonal support for the regulatory programs proposed by FORGE’s statistically weaker single-cell methods.(A) Percentage of tested features reaching FDR < 0.05. The TG-versus-WT contrast returns 340 of 219,519 gene-level tests (RNA, 17 cell types) and 76 of 943,015 peak-level tests (ATAC, 17 cell types); an internal positive control run through identical machinery (microglial/immune versus OPC/oligodendrocyte accessibility, paired within mouse) returns 33,759 of 65,079 peaks. Counts use one label resolution per arm so that no cell contributes twice. Log scale. (B) Significant genes per cell type, cell-level MAST versus replicate-aware edgeR; 17,804 calls become 340. Dashed line, parity. Colour marks the three glial populations carrying the Mef2c program. (C) Program-level testing returns the same answer. Each grey point is one direct activating eRegulon (44-47 per cell type) tested with fry, a rotation-based gene-set test that remains valid at low residual degrees of freedom and with a continuous covariate, and that preserves the observed inter-gene correlation. No regulon reaches directional FDR < 0.05 in any glial population (0 of 47, 0 of 47, 0 of 44); filled diamonds mark Mef2c. The same test on the positive control contrast recovers 44 of 47 regulons at FDR < 0.05, so the TG-versus-WT result reflects power rather than test failure. Open circles show fry’s non-directional (“mixed”) statistic, the single set-level signal that survives: microglial Mef2c targets do respond to condition without a coherent direction (FDR = 0.028), as do 15 of 44 regulons. Dashed line, FDR = 0.05. (D) Log2 fold changes agree. Every gene called by MAST (FDR < 0.05) is plotted against its replicate-aware pseudobulk estimate in the same cell type; the vertical axis is filtered only on testability, never on significance. Pooled Pearson r = 0.61 with 73.0% directional agreement (n = 17,733). Colour by class: glia r = 0.77 pooled (0.82-0.95 per cell type), 90% directional; deep-layer excitatory r = 0.44 (0.36-0.50), 65% directional; other cell types r = 0.90 (0.67-0.97), 96% directional. Only 127 of 17,733 points (0.72%) are themselves pseudobulk-significant. Dashed line, y = x; axes clipped at +/- 5 log2FC with all statistics computed on unclipped values. (E) The same comparison on the ATAC arm, cell-level snapATAC2 against replicate-aware edgeR. Filled points are peaks called in subclasses clearing the max(50, 1% of cells) = 183-cell resolution floor enforced by the pseudobulk build (r = 0.60, 95% directional agreement, n = 112); hollow points are calls from subclasses below that floor (n = 3,031, r = 0.22, 53% directional agreement, i.e. chance), of which 3,028 arise from a single 69-cell subclass for which snapATAC2 returns only 119 distinct log2 fold change values across 3,143 calls. (F) Restricted to the 2,506 genes of the direct activating Mef2c eRegulon, the gene set on which the regulatory claim rests, agreement is undiminished: r = 0.92 and 98% directional in astrocytes (n = 316 MAST-called targets), r = 0.85 and 88% in oligodendrocytes (n = 876), r = 0.93 and 95% in microglia (n = 91), while 0, 0, and 5 of those targets respectively reach pseudobulk FDR < 0.05.

## DISCUSSION

### FORGE corroborates and extends established cell type annotations and regulatory inference in a public PBMC dataset

To establish confidence in FORGE’s outputs, we aligned FORGE’s annotations and regulatory calls with independent analyses of the same public 10x PBMC snMultiome dataset (54,59). We report the confusion matrix from FORGE’s own CellTypist-derived annotations projected against labels transferred onto the same 9,327 cells from the Azimuth human PBMC reference (59) via Seurat reference projection (Figure S1A). Collapsing both vocabularies to lineage, the two agree on 98.4% of cells (T 99.7%, B 98.3%, monocyte 98.1%, NK 87.6%, dendritic 80.2%). At Azimuth’s finer l2 resolution, agreement is ARI 0.661 and AMI 0.675 across 14 CellTypist and 24 Azimuth classes. Residual disagreement is confined to sub-lineage boundaries and is largely due to CellTypist classifiation labels occupying an intermediate granularity between Azimuth’s broad and fine level annotations, rather than systematic misclassification. Additionally, we report a reference-side asymmetry. Azimuth assigns 38.9% of all cells to CD4 TCM while leaving CD4 TEM nearly vacant (64 cells, 0.7%); thus both FORGE Tcm/Naive helper and Tem/Effector helper classes necessarily collapse. Lastly, eight Azimuth classes with no CellTypist counterpart (gdT, dnT, ILC, NK_CD56bright, HSPC, Plasmablast, Platelet, NK Proliferating) are populated but rare.

More critically, we sought to compare and contrast the regulatory programs FORGE surfaces with those recovered by independent methods. During efforts to construct and validate the GLUE software, this same dataset was analyzed via peak-gene link inferecne from the cosine similarity of learned peak and gene embeddings (≤150 kb) (54,60). Comparison with this published analysis simultaneously illustrates concordance and also reveals nuances attributable to underlying method biases inherent to regulatory inference from single-cell multiome data. FORGE robustly recovered the previously reported SPI1-NCF2 relationship in monocytes (61) through two independent regulatory layers, with concordant support from DORC peak-gene associations and SPI1-containing SCENIC+ eRegulons (Figure S1). In contrast, the previously reported lymphoid CD83 program (60) showed only partial concordance (Figure S1). GLUE implicated RELB, BCL11A, and PAX5 in lymphoid-associated CD83 regulation (60), whereas FORGE preferentially assigned CD83 to a myeloid-associated regulatory module containing SPI1, KLF4, MEF2C, CEBPB, BACH1, and LEF1 (Figure S1).

Importantly, the discordance was dependent on the regulatory evidence layer examined rather than representing complete absence of the published lymphoid signal. Depth normalized coverage maps of CD83 demonstrate compareable lymphoid and myeloid accessibility (Figure S1 C). Additionally, at the more permissive DORC peak-gene layer, FORGE recovered RELB as a promoter-proximal relationship exhibiting motif enrichment among CD83-linked regulatory regions; notably, this association did not remain significant after multiple-testing correction (Figure S1). In contrast, BCL11A and PAX5 lacked significant motif enrichment in CD83 DORCs, whereas SPI1 showed significant enrichment after correction (Figure S1). Thus, FORGE retains evidence for a component of the previously reported lymphoid program (60) while providing stronger support for an alternative myeloid-associated regulatory architecture. These differences likely reflect, at least in part, distinct assumptions and evidence requirements of the two frameworks. GLUE incorporates prior regulatory information, including ENCODE-derived TF-binding relationships (60), whereas FORGE’s SCENIC+ and DORC analyses prioritize regulatory associations supported within the measured multiomic dataset and apply different filtering criteria at successive inference layers. Neither framework therefore constitutes ground truth. Rather, the CD83 example illustrates how regulatory conclusions can depend on the evidence model used and highlights the value of exposing multiple independent regulatory layers instead of collapsing them into a single inferred network. Thus, a fruitful future direction may be to incorporate GLUE’s regulatory inference machinery as yet another valueable orthogonal axis of evaluation.

**S1.**
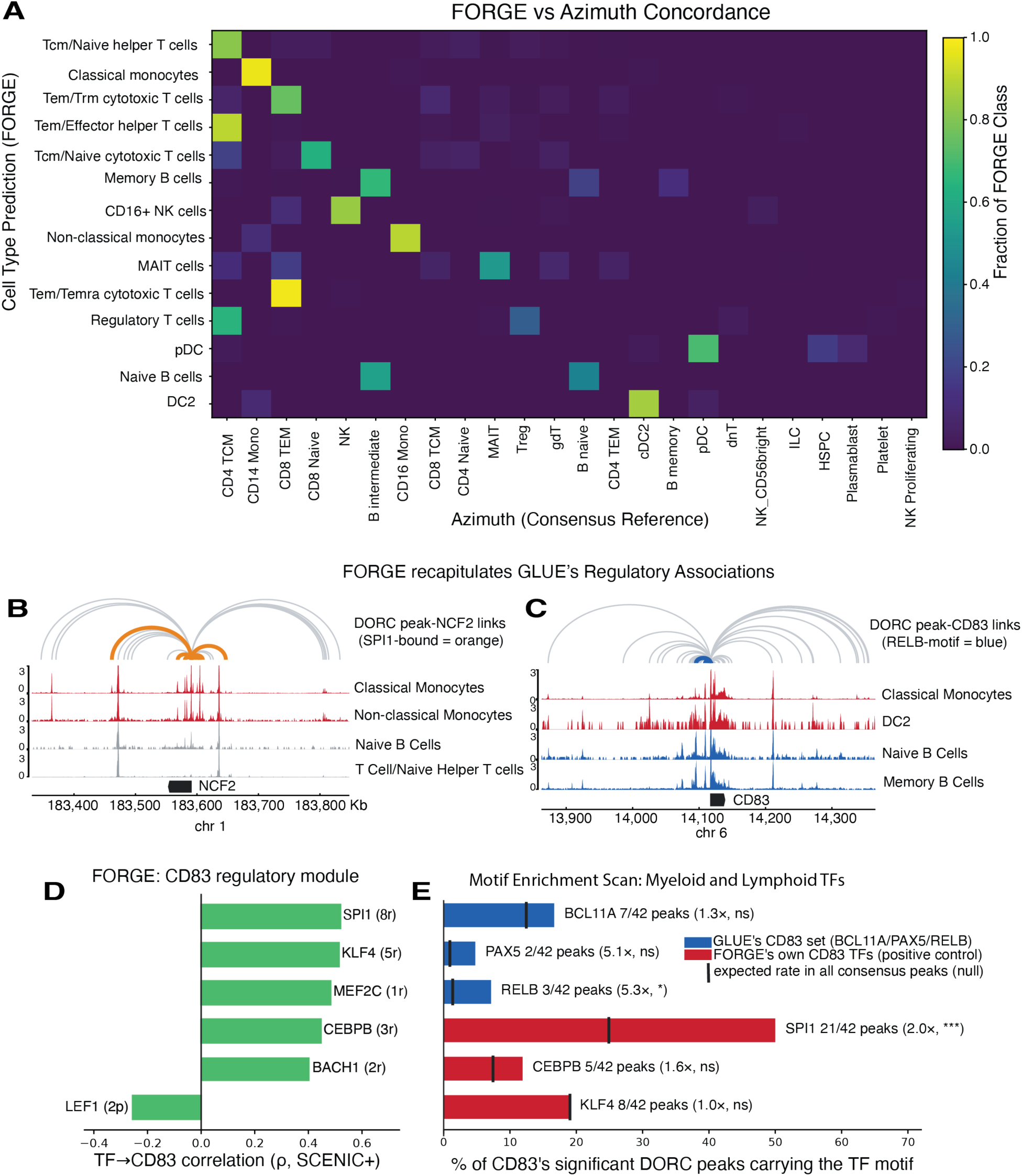
FORGE produces concordant cell-type annotations to Azimuth reference projection at lineage resolution and orthogonally recapitulates regulatory associations identified by GLUE in human PBMC. FORGE excels in biological interpretability when benchmarked against gold-standard tools for integration and generating putative cross modally justified mechanisms of action. (A) Cross-method concordance matrix of 14 FORGE broad CellTypist classes against an inferred reference of labels transferred from the multimodal PBMC reference via Seurat reference projection (FindTransferAnchors/MapQuery) wherein 24 Azimuth l2 classes are observed. The matrix is row-normalized. (ARI=0.661, AMI=0.675, 98.4% agreement when collapsed to the lineage level). Azimuth l2 prediction score mean=0.79 and median=0.85 (11% < 0.5); per-cell mapping score mean=0.63 (30% < 0.5). 10x pbmc_granulocyte_sorted_10k n = 9,327 cells and n=7,620 paired cells. (B) FORGE recapitulates GLUE’s SPI1 to NCF2 regulatory association in monocytes. 31 significant DORC peak-gene links to NCF2; orange = 9 links carrying SPI1 SCENIC+ eRegulon region (12 SPI1 eRegulon regions target NCF2 in total). Depth-normalized ATAC pseudobulk coverage below, with B and T cells as lineage contrasts to the monocyte-biased signal. (C) scGLUE assigns lymphoid TFs whereas FORGE asserts myeloid eRegulons; additionally, FORGE corroborates possible CD83-RELB regulatory action but finds insufficient evidence for scGLUE identified TFs, BCL11A and PAX5. Of the 42 significant DORC links, blue arcs indicate 3 peaks carrying a RELB motif. Depth-normalized ATAC psuedobulk coverage in myeloid and lymphoid cells demonstrates CD83 accessibility in both lineages. (D) FORGE’s CD83 regulatory module (SCENIC+). Bars give ρ, the correlation across cells between each TF’s expression and CD83 expression (rho_TF2G), split at zero by sign; LEF1 is the only negatively correlated TF and is called arepressor by SCENIC+. Parentheses give the number ofdistinct regions (r) linking that TF to CD83. Each is a (TF,region, gene) triplet with three independent lines of evidence: enrichment of the TF’s binding motif in the region, region accessibility vs CD83 expression correlation, and lastly correlation between TF and CD83 expression. None of GLUE’s CD83 TFs (BCL11A, PAX5, RELB) reach CD83 per FORGE’s SCENIC+ implementation. (E) Matched-layer motif test. GLUE’s CD83 call comes from a permissive peak-gene graph, so a more comparable FORGE layer is DORC, not eRegulons. Bars give the percentage of CD83’s 42 significant DORC peaks overlapping (≥1 bp) each TF’s pycisTarget cistrome. Black ticks give that TF’s cistrome as a fraction of all 172,884 consensus peaks. Blue = GLUE’s CD83 set, red = FORGE’s own CD83 TFs (positive control). SPI1 21/42 vs 10.5 expected = 2.0× (one-sided binomial p=3.9e-4, significant after Bonferroni correction across the six TFs tested); RELB 3/42 vs 0.6 expected = 5.3× (uncorrected p=0.019, n.s. after correction); PAX5 2/42 = 5.1× and BCL11A 7/42 = 1.3×, neither significant.

### FORGE Generalizes to Human and Mouse snMultiome Data

In a CRND8 Alzheimer’s disease mouse model 10x snMultiome dataset (55), FORGE QC retained 16,759 paired nuclei across seven broad classes (Figure S2 A-B). Computationally orthogonal MOFA+ and MultiVI embeddings agreed closely (≥0.82 for major cell type classes), with minor discrepancies limited to discernment of vascular from oligodendrocytes and even fewer due to separation of excitatory and inhibitory neurons (Figure S2 C). As in PBMC, rich biological variance is captured via MultiVI and MOFA+ across structured latent spaces (Figure S2 B-E). scPrinter recovered 4,258 DORCs enriched for synaptic and neuronal loci (Figure S2 F), and eRegulon specificity scoring returned canonical lineage regulators (Spi1/Irf8, microglia; Sox10, oligodendrocyte; Dlx1/Lhx6, inhibitory neuron) (Figure S2 G), indicating that FORGE recovers established regulatory architecture without supervision in a second species and tissue.

**Figure S2.**
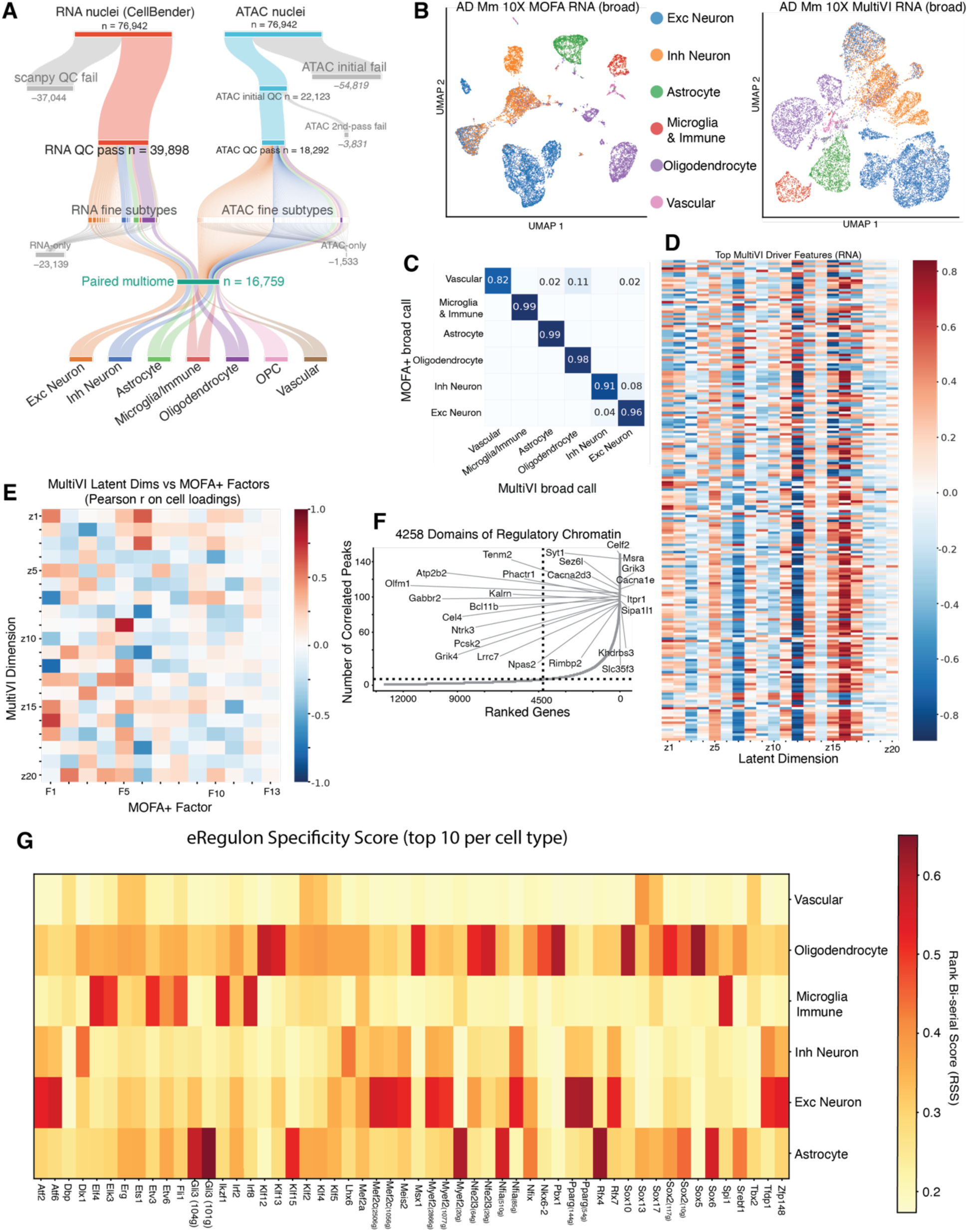
FORGE’s global descriptive architecture scales to multiple samples in a mouse disease model. FORGE end-to-end analysis in an Alzheimer’s disease mouse model, demonstrating QC outputs, integrated dimensional reduction, and gene regulatory network inference in a second species and tissue. (A) Quantitative summary of FORGE QC steps, which retain 39,898 of 76,942 CellBender-denoised RNA nuclei and 18,292 two-pass-filtered ATAC nuclei to identify 16,759 paired multiome nuclei distributed among seven broad cell types. Ribbon width is proportional to nuclei number and modality-restricted nuclei (RNA-only, 23,139; ATAC-only, 1,533) are excluded from downstream analysis. (B) UMAP visualizations of MOFA+ and MultiVI multiome latent embeddings colored by broad RNA annotation, showing that both orthogonal integrations recover the same broad tissue architecture. (C) A KNN classification task assessing agreement between broad cell type annotations within orthogonal latent multiome embeddings, with concordance ≥0.82 for all major classes. (D) Gene expression signatures associated with the top 20 MultiVI dimensions suggest varying levels of the sensitivity of specific MultiVI dimensions to changes in expression. (E) Pearson correlations between MOFA+ factor cell loadings and MultiVI latent embedding dimension cell loadings show emergent agreement among orthogonal methods without one-to-one factor correspondence. (F) 4,258 domains of regulatory chromatin (DORC) orthogonally computed via scPrinter, dominated by synaptic and neuronal loci, corroborate various eRegulon signals; the dotted line marks the correlated-peak threshold and the dashed line the resulting DORC cutoff. (G) Various transcription factors plotted by their respective eRegulon’s rank bi-serial score show oligodendrocytes and excitatory neurons dominate the strongest signal for potential eRegulon-based mechanisms of action, with canonical lineage regulators observed

To evaluate portability across library-preparation technologies, we applied FORGE to a mouse brain snMultiome generated using the BD/Waters platform (Figure S3). Thus, within the two multiome chemistries evaluated here, FORGE processed distinct input conventions while preserving the same downstream analytical framework. Despite the smaller input and differences in preparation quality, QC retained 2,434 paired nuclei resolving into seven broad classes (Figure S3A-B). Broad cell-type structure was concordant between MOFA+ and MultiVI representations, with residual discrepancies concentrated near neuronal subtype boundaries (Figure S3C). scPrinter recovered 6,913 DORCs enriched for synaptic and neuronal loci, and eRegulon specificity scoring recovered canonical lineage regulators (Figure S3G). These results support portability across the tested 10x and BD/Waters datasets without implying universal chemistry independence beyond the platforms evaluated here.

**Figure S3.**
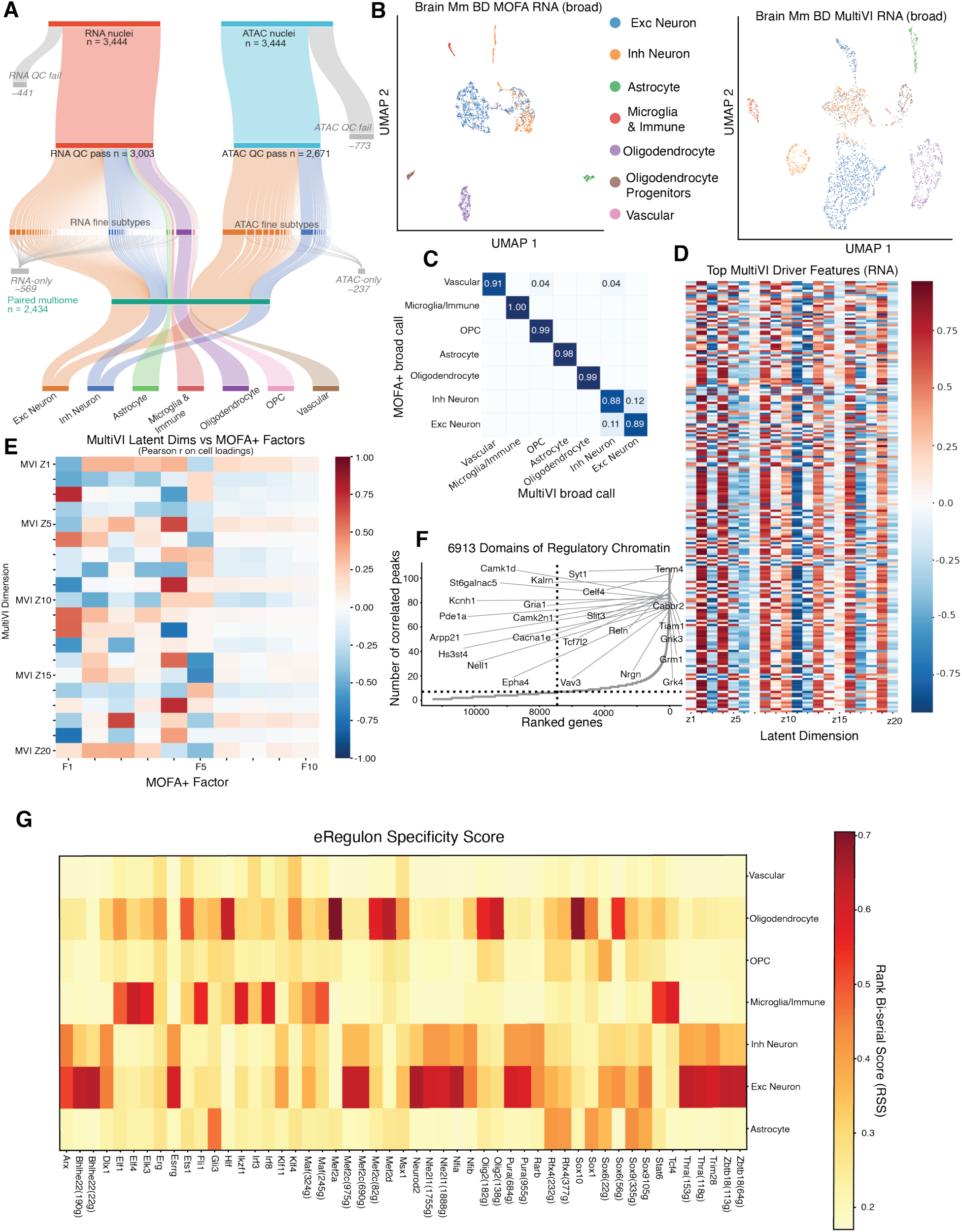
FORGE performs consistently across library preparation chemistries, recovering similar cell type structure and regulatory programs in a BD/Waters mouse brain multiome. FORGE end-to-end analysis in a healthy control mouse brain, demonstrating QC outputs, integrated dimensional reduction, and gene regulatory network inference in a second platform chemistry. (A) Quantitative summary of FORGE QC steps applied to a BD/Waters mouse brain multiome (3,444 nuclei per modality), retaining 3,003 RNA and 2,671 ATAC nuclei and identifying 2,434 paired multiome nuclei distributed among seven broad cell types. Ribbon width is proportional to nuclei number and modality-restricted nuclei (RNA-only, 569; ATAC-only, 237) are excluded from downstream analysis. (B) UMAP visualizations of MOFA+ and MultiVI multiome latent embeddings colored by broad RNA annotation. (C) Excitatory-inhibitory neuron boundary subtyping errors dominate discrepancies in a KNN classification task assessing agreement between broad cell type annotations within orthogonal latent multiome embeddings, with concordance ≥0.98 for all non-neuronal classes. (D) Gene expression signatures associated with the top 20 MultiVI dimensions suggest varying levels of the sensitivity of specific MultiVI dimensions to changes in expression. (E) Pearson correlations between cell loadings within MOFA+ factors (F1-F10) and MultiVI latent embedding dimensions (z1-z20) show emergent agreement among orthogonal methods, concentrated in the leading MOFA+ factors. (F) 6,913 domains of regulatory chromatin (DORC) orthogonally computed via scPrinter corroborate various eRegulon signals; the dotted line marks the correlated-peak threshold and the dashed line the resulting DORC cutoff. (G) Various transcription factors plotted by their respective eRegulon’s rank bi-serial score show oligodendrocytes and excitatory neurons dominate the strongest signal for potential eRegulon-based mechanisms of action, with canonical lineage regulators observed.

FORGE’s capabilities extend to non-immune, non-neural tissue. As a final test of generality, we applied FORGE to a mouse kidney multiome prepared with BD/Waters chemistry, where QC retained 1,556 paired nuclei spanning eight broad classes. Agreement between orthogonal MOFA+ and MultiVI broad calls was strong for transcriptionally distinct compartments (proximal tubule 1.00, vascular 0.97, stromal 0.92) but degraded within the tubule itself, where loop of Henle (0.49) and collecting duct (0.51) nuclei were preferentially reassigned toward distal tubule (0.40 and 0.43, respectively) (Figure S4 C). This pattern mirrors the T cell subtyping discrepancies observed in PBMC and the excitatory-inhibitory boundary in brain; disagreements are due to improper resolution at a granularity finer than the broad ontology. scPrinter recovered 4,302 DORCs enriched for vascular and stromal signaling loci (Flt1, Fli1, Epas1, Ebf1, Eng, Tgfbr3), and eRegulon specificity scoring returned canonical tissue-appropriate regulators (Hnf4a, Esrrg/Esrrb, Pparg, Nfia, Pbx1 in proximal tubule; Fli1, Elk3, Nr2f2 in vascular; Hic1, Klf12 in stromal), confirming that FORGE recovers established regulatory architecture in a third tissue with no meaningful organ-specific tuning.

**Figure S4.**
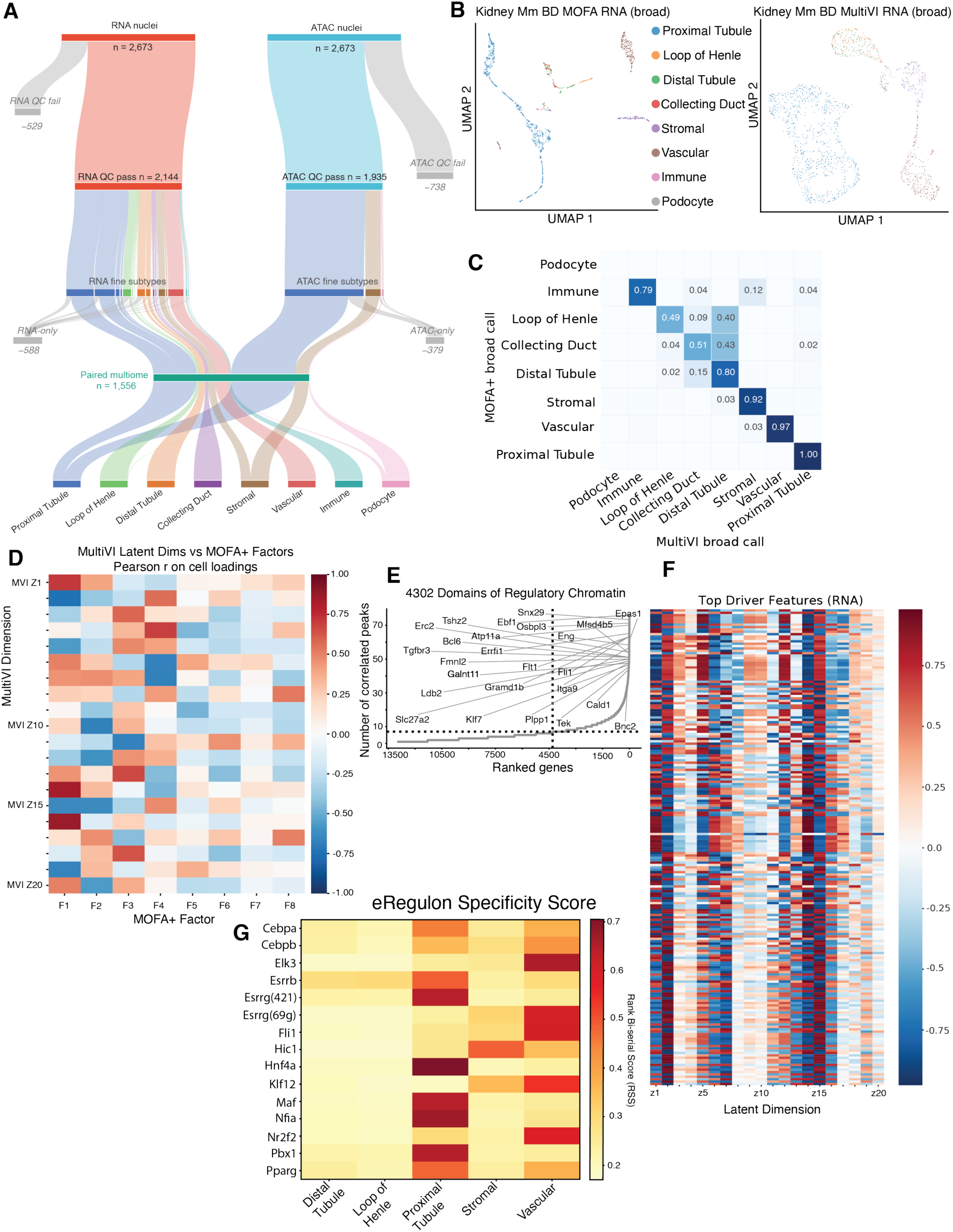
FORGE recovers kidney cell type structure and canonical nephron regulatory programs from a BD/Waters mouse kidney multiome. FORGE end-to-end analysis in a healthy control mouse kidney, demonstrating QC outputs, integrated dimensional reduction, and gene regulatory network inference in a second tissue with an alternative platform chemistry. (A) Quantitative summary of FORGE QC steps applied to a BD/Waters mouse kidney multiome (2,673 nuclei per modality), retaining 2,144 RNA and 1,935 ATAC nuclei and identifying 1,556 paired multiome nuclei distributed among eight broad cell types. Ribbon width is proportional to nuclei number and modality-restricted nuclei (RNA-only, 588; ATAC-only, 379) are excluded from downstream analysis. (B) UMAP visualizations of MOFA+ and MultiVI multiome latent embeddings colored by broad RNA annotation. (C) Loop of Henle and collecting duct subtyping errors dominate discrepancies in a KNN classification task assessing agreement between broad cell type annotations within orthogonal latent multiome embeddings. Observed concordance was ≥0.92 for proximal tubule, vascular, and stromal compartments while misassignment of contiguous tubule segments was biased toward distal tubule populations. (D) Gene expression signatures associated with the top 20 MultiVI dimensions suggest varying levels of the sensitivity of specific MultiVI dimensions to changes in expression. (E) Pearson correlations between MOFA+ factor cell loadings (F1-F8) and MultiVI latent embedding dimension cell loadings (z1-z20) show emergent agreement among orthogonal methods and illuminate opportunities for exploration where anti-correlations indicate non-linear biology. (F) 4,302 domains of regulatory chromatin (DORC) (≥7 correlated peaks) orthogonally computed via scPrinter corroborate various eRegulon signals; the dotted line marks the correlated-peak threshold and the dashed line the resulting DORC cutoff. (G) Various transcription factorss plotted by their respective eRegulon’s rank bi-serial score show proximal tubule and vascular compartments dominate the strongest signal for potential eRegulon-based mechanisms of action; parenthetical values indicate the number of target genes per eRegulon. Cell types with insufficient nuclei for stable eRegulon scoring (podocyte, immune, collecting duct) are not shown.

### Parallel evaluation of cell- and sample-level differential analyses defends Mef2c regulatory program viability

The juxtaposed merits of replicate-aware pseudobulk aggregation and cell-level testing remain contested. The arguments trend towards aggregation of biological replicates as a necessary prerequisite for valid inference; however, cell-level tests invariably retain information that aggregation discards as noise (62,63). We report both analyses and let them answer different questions. The statistically grounded replicate- and age-aware tests scope what this cohort can establish inferentially, and by that standard the condition effects do not survive rigorous corrections (Figures 4A-4C) for individual genes, peaks, nor whole eRegulon programs. We attribute the apparent discrepancy to a six-animals-per-group design rather than evidence against the underlying biology. To orthogonally query potential agreement in underlying signals between the methods, we additionally considered the concordance of effect sizes. The two analyses differ in unit of replication, noise model, and software, so the agreement of their log2 fold changes on the Mef2c target set itself (Figure 4D-4F) support that the direction and relative magnitude of changes are not type 1 artifacts. In fact, cell-type dependence surfaces to further bound where we place confidence: concordance is high in the glial populations carrying the Mef2c program and close to chance in deep-layer excitatory neurons (Figure 4D-4E). Thus, we may scope our regulatory claims to the former which originally received support from the pseudoreplicate analysis interpretation. We present the Mef2c-centred regulatory relationships as hypotheses supported by convergent computational evidence (motif enrichment, footprint occupancy, eRegulon activity, and reproducibility of effect size under orthogonal replication) (Figures 3, 4D-4F) rather than as conclusive findings. A natural next step is experimental validation in an appropriately powered cohort and targeted perturbation of Mef2c and predicted target enhancers.

### Validation Summary and Reconstructive Masking Sweep Quantifications

Thus, to establish that FORGE generalizes beyond any single experimental context, we assembled a validation set spanning four independent snMultiome datasets (54,55) that vary along four essential axes: species (h. sapiens, mus musculus), library chemistry (10x Genomics, BD/Waters), tissue (blood, brain, kidney), and experimental design (single-sample, comparative multi-sample) (Figure S5). Human PBMC on 10x chemistry demonstrates functionality in high-quality human cell-types with an established dataset. Similarly, the CRND8 mouse cortex cohort (12 samples, transgenic versus wild-type) critically exercises condition-aware multisample scaling in a disease model (Figure S5). The BD/Waters brain and kidney datasets indicate the architectures’ adaptability to platform chemistry and tissue sources.

**Figure S5.**
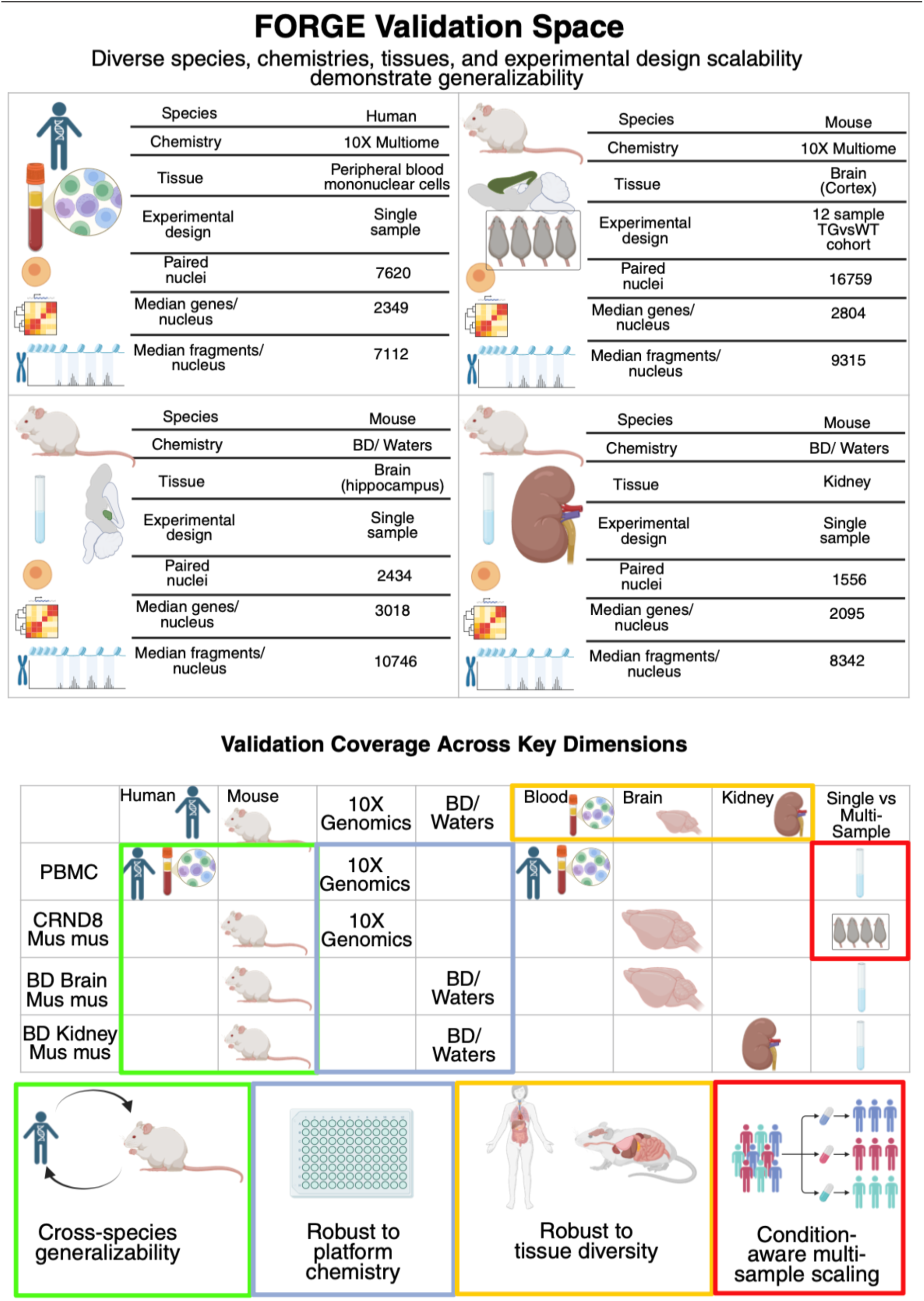
The FORGE validation space comprising diverse species, platform chemistries, tissues, and experimental designs demonstrates generalizeability. Summary characteristics of the four snMultiome datasets used to validate FORGE. For each dataset, species, library chemistry, tissue of origin, experimental design, number of paired nuclei passing FORGE quality control, and median genes and ATAC fragments recovered per nucleus are reported: human PBMC (10x Multiome, single sample), mouse cortex (10x Multiome, 12-sample transgenic versus wild-type cohort), mouse hippocampus (BD/Waters, single sample), and mouse kidney (BD/Waters, single sample). Coverage matrix indicating which validation dimension each dataset addresses. Rows are datasets; columns are species, chemistry, tissue, and experimental design. Cross-species generalizability (green; human PBMC versus mouse datasets), robustness to platform chemistry (blue; 10x versus BD/Waters within mouse), robustness to tissue diversity (yellow; blood, brain, and kidney), and condition-aware multisample scaling (red; single-sample designs versus the 12-sample CRND8 cohort).

MultiVI’s original report established that the model can impute a withheld modality for cells observed in only one assay (39,50). We reproduced that experiment inside FORGE. For human PBMC (54), mouse brain, and mouse kidney (BD/Waters), new MultiVI models were trained with 25%, 50% and 75% of cells artificially stripped of one modality and the withheld values scored against measured known values: per-cell Spearman ρ for withheld RNA, per-cell AUPRC for withheld ATAC (n masked cells per sweep iteration; Figure S6A). Reconstruction fidelity is flat across the masking gradient in all three datasets (Figure S6B) with joint-latent quality preserved (Figure S6C). Absolute fidelity also closely mirrors the MultiVI publication values (39,50) (carets, Figure S6A).

**Figure S6.**
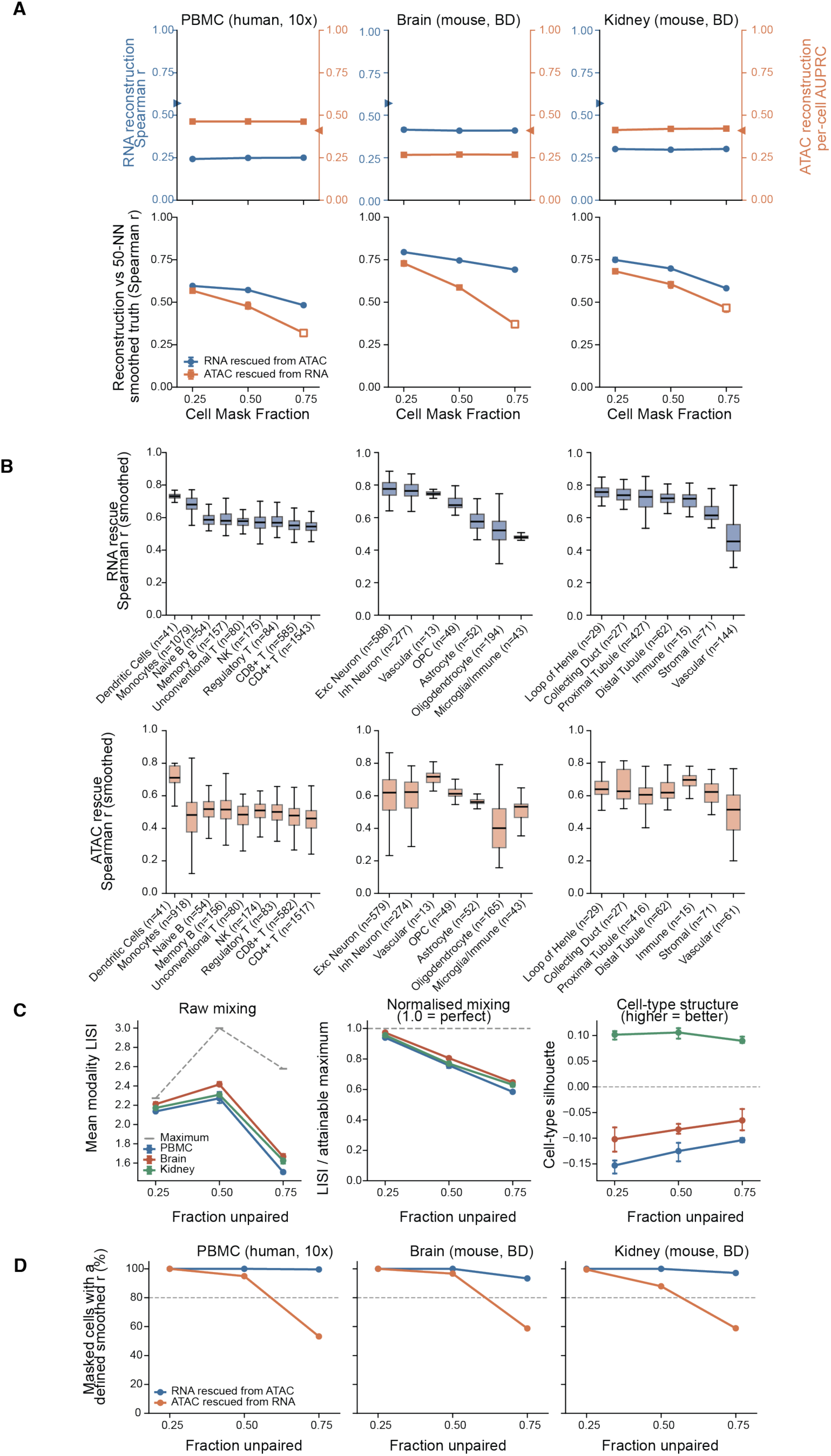
Cross-modal reconstruction under artificial unpairing. Across three tissues, two species and two chemistries, FORGE’s MultiVI integration reconstructs a withheld modality with fidelity that is statistically indistinguishable at 25%, 50% and 75% artificial unpairing (A) Top Row: Reconstruction of a withheld modality as a function of the fraction of cells artificially masked, for three FORGE datasets (columns). Y-axes illustrate per-cell Spearman correlation of imputed versus true expression for cells denied RNA (blue, left axis, circles) and per-cell AUPRC of imputed versus true accessibility for cells denied ATAC (orange, right axis, squares). The two axes span the same 0-1 range and neither is rescaled, though they carry different metrics whose absolute levels are not comparable; thus the panel should be read for slope within each series independently. Carets on each spine mark the corresponding MultiVI-publication benchmark. Bottom Row: both directions scored against a 50-nearest-neighbour smoothed target, which is a common scale. Points are the mean of three per-seed medians. Hollow markers flag points where the metric was undefined for >20% of masked cells (see D). (B) Reconstruction by broad cell class at 50% masking, pooled over seeds; boxes show median and IQR, whiskers 1.5 x IQR, ordered by RNA-rescue median with the ATAC row held inthe same order; n is masked cells per seed. (C) Joint-latent quality. Left: raw modality LISI against its composition-dependent attainable maximum (dashed). Middle: the same score normalised by that maximum. Right: cell-type silhouette. (D) Percentage of masked cells for which the smoothed correlation in (A, bottom row) is defined; the smoothed target is built from the masked object, so it becomes uninformative for ATAC at high unpairing. Dashed line, 80%.

### Limitations of the study

FORGE was designed and evaluated in human and mouse datasets from blood, brain, and kidney and across two multiome chemistries. Performance in other organisms, tissues, assay chemistries, and substantially different data-quality regimes remains to be established. Dynamic QC thresholds reduce reliance on fixed cutoffs but may require tissue-specific review, particularly for cell populations with unusual RNA content or chromatin profiles. Several differential modes operate at the cell level and are best interpreted as exploratory state comparisons when biological replicates are available (64,65). Critically, given only male samples were evaluated, the datasets used to validate FORGE make it impossible to assess whether FORGE can robustly identify sex-differences which are simultaneously known to be under-characterized as well as bear significant biology. Moreover, we must also consider analyses ingesting DEGs produced via cell-level psuedoreplicate approaches. For example, while we discuss the putative Mef2c regulatory program at length and provide strong justification for future work to empirically verify our findings, it would not be entirely suprising if a properly powered lab experiment ultimately did not yield support for such a regulatory system. Additionally, differential module eigengene testing in hdWGCNA remains a cell-level Wilcoxon rank-sum test, whereas the differential expression and accessibility results reported here are tested at the sample level. Module-trait and DME contrasts are therefore subject to the same inflation of effective sample size that motivated the pseudobulk framework, and we read them as descriptive rather than as replicate-aware evidence. Metacell construction grouped by cell type and sample absorbs part of the within-sample correlation, but a sample-level test of module eigengenes would be the stricter analysis and we have not performed one. Most importantly, our reported findings are a more accurate reflection of the capabilities and limitations of FORGE’s comprising toolkit rather than a reflection of incorrect or inappropriate architecture mapping the various tool inputs and outputs to each other.

Two asymmetries limit the comparison of FORGE’s FOSCTTM against the published GLUE baseline. FOSCTTM is a fraction and is therefore normalized by pool size, but as cell density rises a fixed embedding displacement is out-ranked by more cells, so values computed on pools of different size are not directly comparable; FOSCTTM recomputed on a subsample matched to the GLUE cell count is reported alongside. More importantly, FORGE’s MultiVI is a paired model, trained on cells whose RNA-ATAC correspondence is known, whereas the GLUE baseline methods perform diagonal integration without pairing. FORGE’s lower FOSCTTM therefore reflects an easier problem and is not evidence of a superior integration method; this asymmetry is characterized directly by reconstruction fidelity under a masking sweep and by FOSCTTM recomputed on artificially unpaired models. Finally, FORGE integrates multiple computational evidence layers to nominate regulatory hypotheses, but motif occurrence, co-accessibility, footprinting, DORCs, eRegulons, and cross-modal concordance do not by themselves establish causal regulatory mechanisms. Experimental perturbation remains necessary for causal validation.

## RESOURCE AVAILABILITY

### Lead contact

Requests for further information and resources should be directed to and will be fulfilled by the lead contact, Vivek Swarup.

### Materials availability

Each 10x Genomics dataset used for validation of FORGE should be referenced for their own materials availability, respectively.

While BD/Waters Bioscience generously supplied Brain and Kidney snMultiome datasets, these materials must be requested from the vendor as the authors do not have authority over distribution.

### Data and code availability

• The Waters Biosciences, formerly BD Rhapsody, data reported in this study cannot be deposited in a public repository because it remains proprietary and only available upon request. To request access, contact the vendor.

• This paper analyzes existing, publicly available data, accessible at “PBMCs from C57BL/6 mice (v1, 150×150), Single Cell Immune Profiling Dataset by Cell Ranger v3.1.0, 10x Genomics, (2019, July 24)”.

• This paper analyzes existing, publicly available data, accessible at “10x Genomics Multiomic Integration Neuroscience Application Note: Single Cell Multiome RNA + ATAC Alzheimer’s Disease Mouse Model Brain Coronal Sections from One Hemisphere Over a Time Course”.

**Instructions for section 1: Data**

• All original data has been deposited at github.com/swaruplabUCI/FORGE and is publicly available as of the date of publication.

**Instructions for section 2: Code**

• All original code has been deposited at github.com/swaruplabUCI/FORGE and is publicly available as of the date of publication.

**Instructions for section 3: Additional information**

• Any additional information required to reanalyze the data reported in this paper is available from the lead contact upon request.

## ACKNOWLEDGMENTS

We thank members of the Swarup laboratory for their helpful comments on the manuscript, and their methodological guidance. We also thank the Research Cyberinfrastructure Center (RCIC) at the University of California, Irvine for providing high-performance computing resources and technical support. This work was supported by the Alzheimer’s Association Research Grant (AARG-25-1472103), the Cure Alzheimer’s Fund, and the National Institute on Aging of the National Institutes of Health under award numbers R01AG071683 and U54AG054349 to V.S.

## AUTHOR CONTRIBUTIONS

Conceptualization, VS and LES; methodology, LES, NR, ZS, and VS; Investigation, LES; writing - original draft, LES; writing - review & editing, LES, NR, ZS, and VS.; funding acquisition, VS; resources, VS; supervision, VS.

## DECLARATION OF INTERESTS

We have no conflicts of interest to declare.

## DECLARATION OF GENERATIVE AI AND AI-ASSISTED TECHNOLOGIES IN THE WRITING PROCESS

During the preparation of this work, the authors used Anthropic’s Claude Opus 4.7 to review coding syntax and coordinate inputs and outputs between various software. Additionally, a draft template for the methods was generated from the repository structure prior to major edits for accuracy and flow. After using this tool or service, the authors reviewed and edited the content as needed and take full responsibility for the content of the publication.

## SUPPLEMENTAL INFORMATION

**Document S1. Figures S1-S6 & Tables S1-S3**

## STAR★METHODS

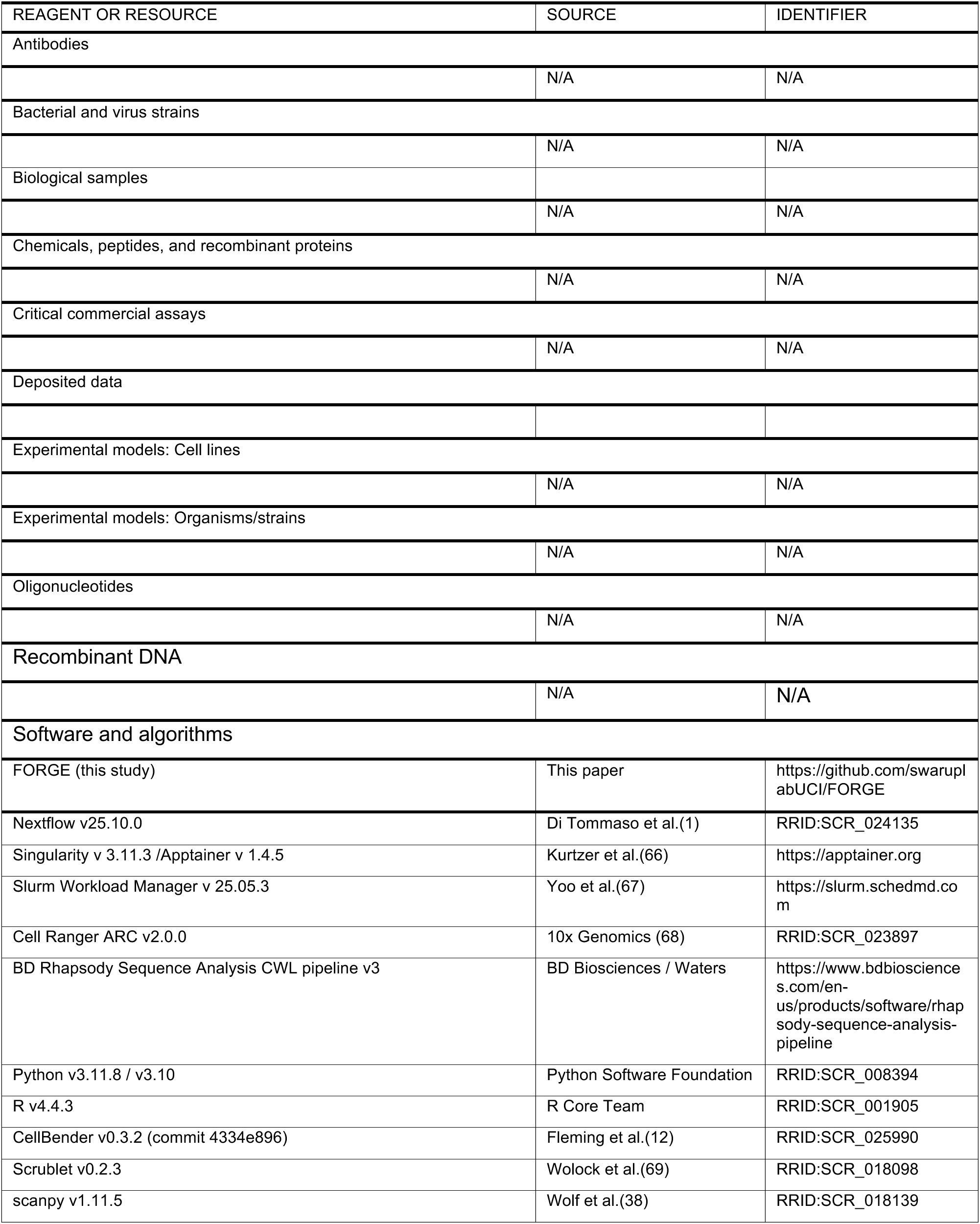
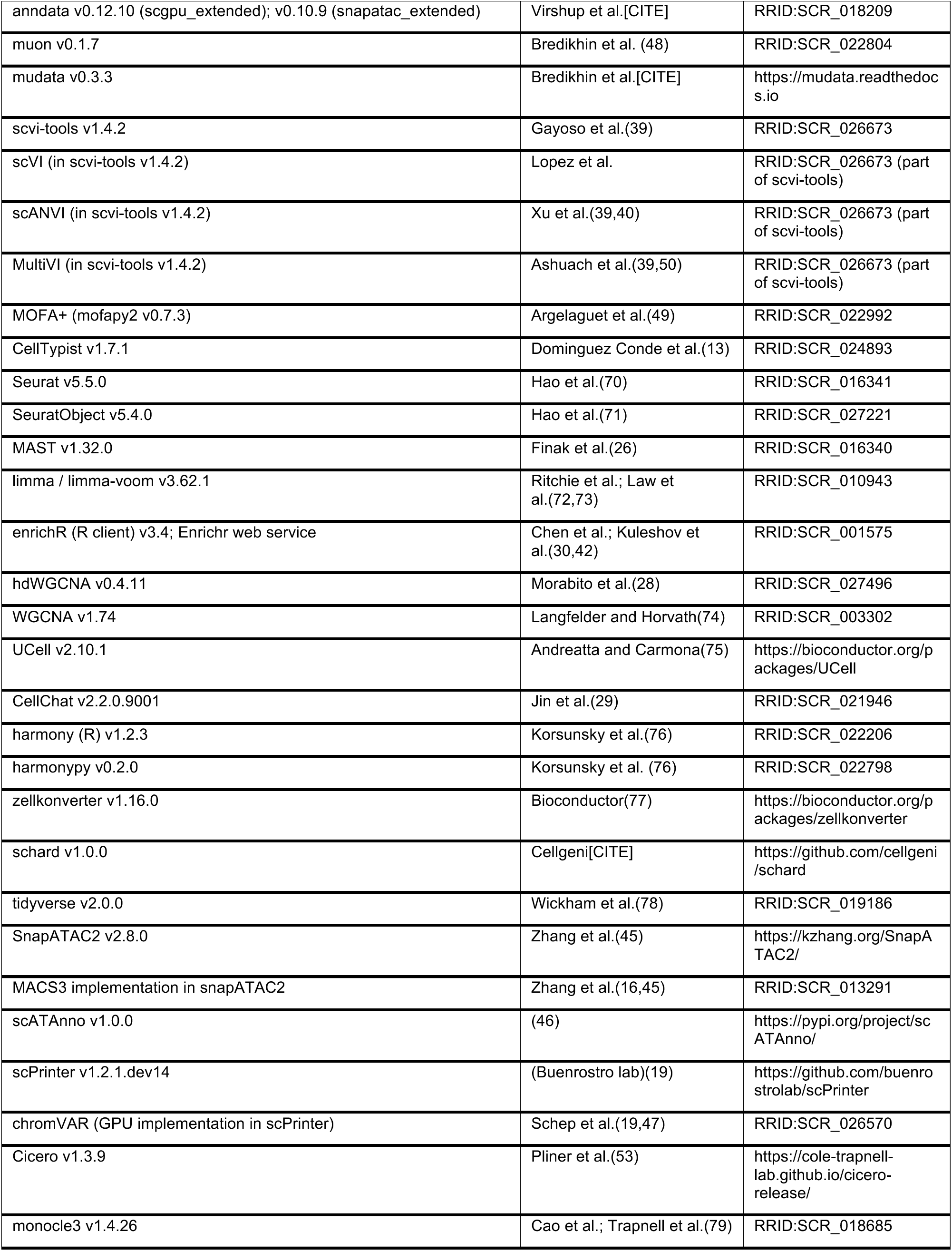
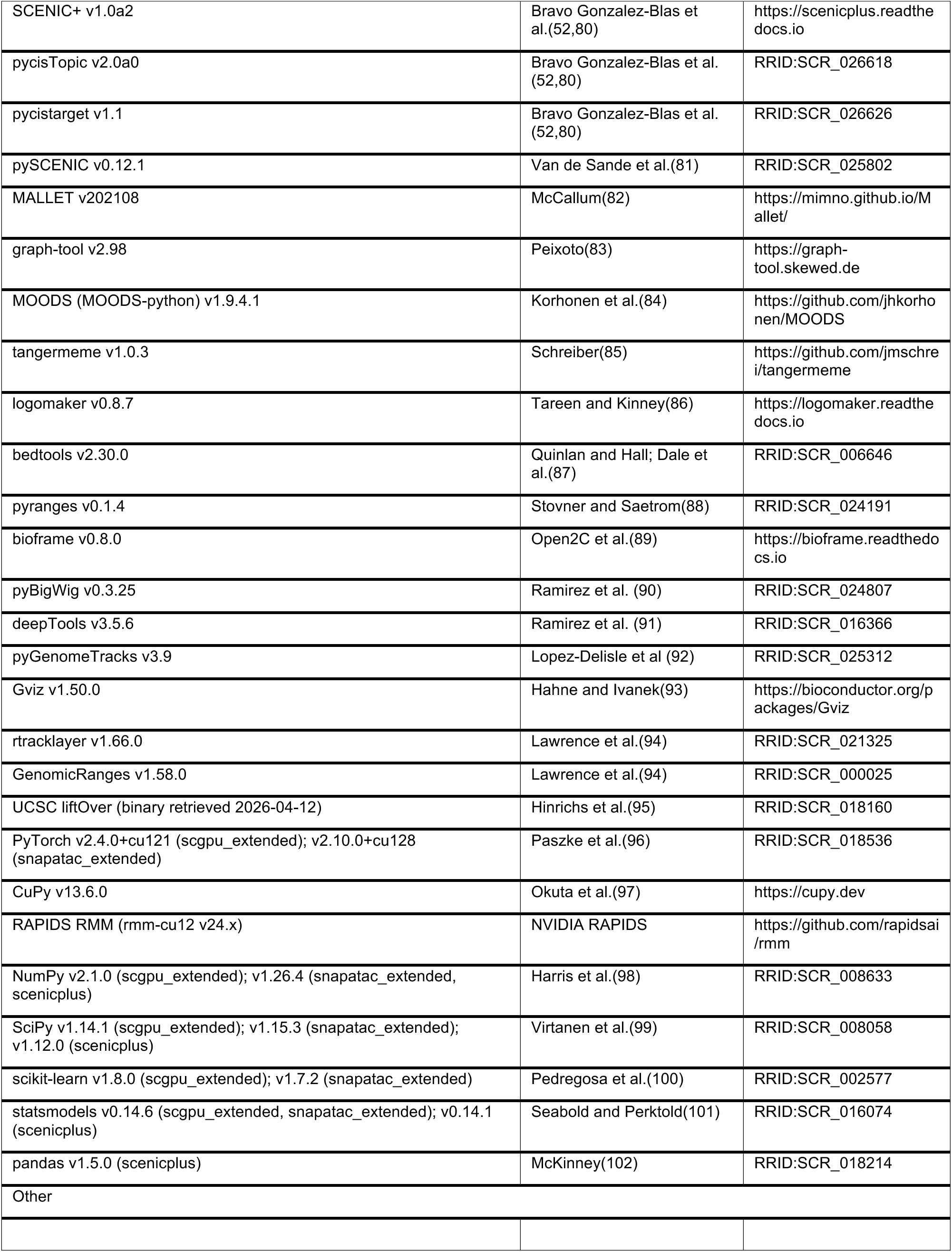
KEY RESOURCES TABLE.

## EXPERIMENTAL MODEL AND STUDY PARTICIPANT DETAILS

Secondary use of existing datasets. This study generated no primary data. All four multiome datasets analyzed here were pre-existing: two are publicly released 10x Genomics demonstration datasets, and two were provided by the BD Rhapsody/Waters Corporation vendor. No animals were bred, housed, treated, or sacrificed by the authors, and no human participants were recruited or sampled by the authors. Secondary computational analysis of these de-identified, previously collected datasets did not constitute human-subjects research or animal research at UCI, and no additional institutional approval was required. Where the originating source reports species, strain, genotype, age, sex, husbandry, or ethical approval, those details are reproduced below and attributed; where the source does not report a field, this is stated explicitly.

I. Human peripheral blood mononuclear cells (10x Genomics public dataset). Species: Homo sapiens. Sample: peripheral blood mononuclear cells, granulocytes removed by cell sorting, from one healthy donor. Age: not reported. Sex: Male. Ancestry, race, ethnicity: not reported by the source. Health status: Healthy. Assay: Chromium Single Cell Multiome ATAC + Gene Expression, [chemistry version], Chromium X instrument; processed by the source with Cell Ranger ARC 2.0.0 against GRCh38. Donor consent and ethical oversight were the responsibility of 10x Genomics and its tissue supplier; the dataset is distributed publicly at [URL]. The authors had no access to identifiable donor information.
II. Mouse Alzheimer’s disease model brain (10x Genomics public dataset). Species: Mus musculus. Strain/model: B6C3-Tg(PRNP-APPSweInd)8Dwst (commonly referred to as TgCRND8), with non-transgenic wild-type littermate controls. Genotype groups: Transgenic (TgCRND8) and wild-type control (exact sample sizes per group not reported in dataset metadata). Age/developmental stage: 2.5, 5.7, 13.2, and 17.9 months of age. Sex: Male. Tissue: brain, coronal sections from one hemisphere. Number of animals and number of sections per animal: 12 animals, 5 µm sections per animal. Assay: Chromium Single Cell Multiome ATAC + Gene Expression; processed by the source against [build]. Animal husbandry, care, and institutional approval were the responsibility of 10x Genomics and its tissue supplier as described in the 10x Genomics Multiomic Integration Neuroscience Application Note.
III. Mouse brain (vendor-provided dataset). Species: Mus musculus. Strain: Vendor unable to disclose. Genotype: wild type. Age: Vendor unable to disclose. Sex: Male. Tissue: fresh snap frozen brain. Number of animals: 1. Tissues were dissociated following the demonstrated protocol: “Nuclei Isolation and Cleanup from Frozen Mouse Brain for Single Nuclei Sequencing Applications” (100-272-080), using the NIC+ Cartridge (100-215-389), RNase Inhibitor V2 (100-288-916), Nuclei Isolation Reagent (100-063-396), Nuclei Storage Reagent (100-063-405), and Nuclei Debris Removal Stock Reagent (100-253-628). Assay: BD Rhapsody™ ATAC-Seq Assay (Kit Cat. No. 571201, 571361) following BD Rhapsody™ Single-Cell ATAC-Seq and mRNA Whole Transcriptome Analysis Library Preparation Protocol (23-24474(02)). The resulting ATAC and WTA libraries were sequenced on the AVITI™ system. BD Cellismo™ 2.0; preprocessed with the vendor CWL workflow. These data were provided by BD Biosciences / Waters Corporation and are not redistributable by the authors, but may be redistributable by the vendor upon request; see Data and code availability. Animal care and institutional approval were the responsibility of the provider.
IV. Mouse kidney (vendor-provided dataset). Species: Mus musculus. Strain: Vendor unable to disclose. Genotype: wild type. Age: Vendor unable to disclose. Sex: Male. Tissue: fresh snap frozen kidney. Number of animals: 1. Tissues were dissociated following the demonstrated protocol: “Nuclei Isolation and Cleanup from Frozen Mouse Brain for Single Nuclei Sequencing Applications” (100-272-080), using the NIC+ Cartridge (100-215-389), RNase Inhibitor V2 (100-288-916), Nuclei Isolation Reagent (100-063-396), Nuclei Storage Reagent (100-063-405), and Nuclei Debris Removal Stock Reagent (100-253-628). For kidney nuclei, the demonstrated protocol was modified by replacing the myelin removal step with one additional wash. These data were provided by BD Biosciences / Waters Corporation and are not redistributable by the authors, but may be redistributable by the vendor upon request; see Data and code availability. Animal care and institutional approval were the responsibility of the provider.

3 mL of Nuclei Storage Reagent supplemented with RNase Inhibitor V2 at 0.2 U/mL .Assay: BD Rhapsody™ ATAC-Seq Assay (Kit Cat. No. 571201, 571361) following BD Rhapsody™ Single-Cell ATAC-Seq and mRNA Whole Transcriptome Analysis Library Preparation Protocol (23-24474(02)). The resulting ATAC and WTA libraries were sequenced on the AVITI™ system. BD Cellismo™ 2.0; preprocessed with the vendor CWL workflow [version]. These data were provided by BD Biosciences / Waters Corporation and are not redistributable by the authors, but may be redistributable by the vendor upon request; see Data and code availability. Animal care and institutional approval were the responsibility of the provider.

## METHOD DETAILS

Equations are given where FORGE defines a quantity, departs from a package’s documented behavior, or where a parameter interaction affects interpretation; published methods used at their documented settings are specified by function and parameters and cited to their original description.

### Pipeline implementation

FORGE is a workflow for end-to-end analysis of paired single-nucleus multiome (snRNA-seq + snATAC-seq) data, implemented as a Nextflow pipeline. The main workflow file declares 129 process modules and path-resolving helper functions, implements a pre-flight validation checklist, and defines twelve parameter-gated sub-workflow blocks together with the default entry points; FORGE architecture and internal organization is documented at the project site given in “additional resources”.

The thirteen sub-workflows and their activation gates are: RNA (rna.run), RNA_DIFFERENTIAL (differential_rna.run), ATAC_INITIAL (atac.run), ATAC_FINAL (atac.run), ATAC_DIFFERENTIAL (differential.run), REGULATORY_ANALYSIS (cicero.run/chromvar.run/scprinter.run), MULTIOME_INTEGRATION (run_multiome_integration), MULTIOME_GRN (pycistopic.run/scenicplus.run), ENHANCER_FOOTPRINTING_RECIPES (enhancer_footprinting.run), SHI_FIGURES (shi_figures.enabled), and VIZ_ONLY (invoked with -entry VIZ_ONLY). Dependency order in the default entry workflow is RNA to RNA_DIFFERENTIAL; ATAC_INITIAL to ATAC_FINAL to {ATAC_DIFFERENTIAL, REGULATORY_ANALYSIS}; {RNA, ATAC_FINAL} to {MULTIOME_INTEGRATION, MULTIOME_GRN}; and {REGULATORY_ANALYSIS, MULTIOME_GRN} to ENHANCER_FOOTPRINTING_RECIPES. Downstream processes require specific upstream outputs to launch. A change to an intermediate process preserves completed upstream work and only invalidates compute downstream of the change.

#### Per-cell-type resolution floor

Every stage that fans out per cell type enforces a common floor: a cell type is analyzed only if it contains at least max(*min*_*cells*, *min*_*pct* × *N*_total_) cells, with defaults *min*_*cells* = 50 and *min*_*pct* = 0.01. This is both a statistical and an operational constraint. Cell types failing the floor are skipped and logged rather than silently dropped. Individual analyses impose additional floors where their own requirements are stricter, and these are collected in Table S2.

### Input specification

All input is declared in a single manifest CSV with columns sample_id, batch, sample_type, original_lane_id, rna_file, fragment_file, condition_group, and data_dir. Rows are parsed, whitespace is trimmed by field, fields are filtered for non-empty contents, and the results are mapped to (sample_id, file) tuples. Path-resolving helper functions maintain file provenance and raise errors on unexpected input. The condition_group column supplies the experimental design axis consumed by the differential comparison modules.

### Execution environment and containerization

Every process executes inside one of five Singularity images (Table S1), each built from a customizable definition file in the repository that specifies the base image, pinned package versions, and any build-time patches; software versions are given in the key resources table. The nextflow.config files and profiles bind these images to the workflow. The withName blocks attach per-process settings and profiles are named configuration bundles selected at launch that swap the execution backend; that is partition, account, quality of service, job concurrency, and container resource labels are easily configurable without editing code. Profile-level withName blocks override top-level withName blocks by design.

Images are run with --contain --home /tmp and explicit bind mounts, with cache directories (Numba, Matplotlib, XDG, CuPy) redirected to /tmp, PYTHONNOUSERSITE=1, and HDF5_USE_FILE_LOCKING=FALSE; R-based processes additionally set R_LIBS_USER=/dev/null to prevent host library leakage. Processes designated GPU-capable (TRAIN_SCVI, TRAIN_SCANVI, CELLBENDER, GPU_CHROMVAR, MOFA_INTEGRATE, MULTIVI_INTEGRATE) request --nv and one accelerator by default. Of these, CELLBENDER and GPU_CHROMVAR require a GPU; the scvi-tools models fall back to CPU when params.scvi_accelerator is set to cpu, at a substantial cost in runtime, and a GPU is recommended.

Resource requests are declared as labels, from process_small (8 GB, 1 CPU) through hugemem (500 GB, 4 CPU), and are specialized per process by a resource tier selected with params.resource_tier (test, tutorial, small, medium, large). All work reported here was executed on the UCI HPC3 SLURM cluster.

**Table S1.** Container images and their principal packages.

| Image | Base image | Principal packages |
| --- | --- | --- |
| scgpu_extended.sif | scverse/scvi-tools:py3.11-cu12-base | scvi-tools, CellBender (pinned commit 4334e89), CellTypist, scanpy, anndata, muon, mudata, mofapy2, Scrublet, PyTorch, NumPy, SciPy, scikit-learn, statsmodels |
| snapatac_extended.sif | python:3.10-slim | SnapATAC2, MACS3, scPrinter, scATAnno, harmonypy, PyTorch, CuPy, RAPIDS RMM, NumPy, SciPy, scikit-learn, statsmodels, anndata, scanpy, pyBigWig, pyGenomeTracks, logomaker, tangermeme, MOODS, pyranges, bioframe, deepTools |
| scenicplus.sif | python:3.11.8-slim | SCENIC+, pycisTopic, pycistarget, pySCENIC, MALLET, bedtools, graph-tool (isolated conda environment), NumPy, SciPy, pandas, statsmodels |
| seurat_extended.sif | rocker/r-ver:4.4.3 | Seurat, SeuratObject, MAST, edgeR, limma, statmod, locfit, hdWGCNA, WGCNA, CellChat, enrichR, UCell, harmony, tidyverse, zellkonverter, schard |
| cicero.sif | condaforge/mambaforge | Cicero, monocle3, Gviz, rtracklayer, GenomicRanges |

**Table S2.** Minimum cell and group sizes by analysis.

| Analysis | Floor | Unit |
| --- | --- | --- |
| All per-cell-type fan-out (default) | max(50, 0.01N) | cells per cell type |
| Differential accessibility, cell-level | 50 | cells per condition group |
| Differential TF accessibility | 50 | cells per condition group |
| chromVAR permutation contrast | 10 (flagged below this) | cells per group |
| Pseudobulk column occupancy | 10 | cells per (cell type, sample) |
| Pseudobulk replication | 3 | retained columns per condition |
| Co-expression networks | 100 cells; 20 metacells | cells, then metacells per cell type |
| Cicero per cell type x condition | 250 | cells per stratum |
| Cell-cell communication | 10 | cells per sender or receiver population |
| Enhancer program decomposition | 200 | cells per group |
| Coverage track export | 100 | cells per group |

### Reproducibility, provenance, and random seeds

Where a method’s own randomness affects the result, seeds are fixed. params.random_seed = 42 is threaded into MOFA+, scVI, scANVI, MultiVI, hdWGCNA, Cicero, and the pseudobulk permutation and subsampling procedures; Leiden clustering uses random_state = 0; the MultiVI masking sweep replicates at seeds 42, 123, and 456; MOFA+ factor stability is bootstrapped over 10 iterations at 50% cell sub-sampling; and LDA sets random state =555.

Every FORGE run emits a Nextflow timeline, an execution report, a directed acyclic graph, and a per-task trace under outdir/pipeline_info/. These are the provenance records used for the compute accounting reported in Figure 1. FORGE additionally supports resumption at defined checkpoints through an on-ramp mechanism (params.onramp), which accepts pre-computed integrated RNA objects, ATAC peak matrices, MuData objects, and scPrinter printer objects.

### Datasets

FORGE was validated on four independent snMultiome datasets spanning two species, two assay chemistries, and both single-condition and multi-sample differential experimental designs (Figure S1) (Table S3). BD Rhapsody datasets were pre-processed through the vendor CWL workflow prior to FORGE; 10x datasets were ingested from raw feature-barcode matrices (raw_feature_bc_matrix.h5) and ATAC fragment files generated by Cell Ranger ARC.

Per-dataset quality-control parameters were set from tissue characteristics rather than applied uniformly: mitochondrial-content ceilings of 5% for mouse brain nuclei (AD, Brain) and 20% for PBMC and kidney; minimum genes per cell of 100 for nuclei and 200 for PBMC; and CellBender expected_cells matched to expected recovery per dataset (8,000 AD; 10,000 PBMC; 3,000 Brain; 2,000 Kidney).

**Table S3.** Validation datasets.

| Dataset | Species | Platform | Samples | Design | Reference build |
| --- | --- | --- | --- | --- | --- |
| PBMC<br>(pbmc_granulocyte_sorted_10k) | Human | 10x<br>Chromium<br>X | 1 | Single<br>healthy<br>donor | GRCh38,<br>GENCODE<br>v38, ENCODE<br>hg38 blacklist<br>v2 |
| AD (TgCRND8 model) | Mouse | 10x<br>Multiome | 12 | Differential,<br>TG versus<br>WT | mm10,<br>GENCODE<br>vM10, mm10<br>blacklist v2 |
| Brain | Mouse | BD<br>Rhapsody<br>multiome | 1 | Single<br>control | GRCm39,<br>GENCODE<br>vM37 |
| Kidney | Mouse | BD<br>Rhapsody<br>multiome | 1 | Single<br>control | GRCm39,<br>GENCODE<br>vM37 |

The AD cohort comprises six transgenic and six wild-type CRND8 mice, assigned from the source sample manifest. It is an age series wherein transgenic and wild-type ages are not fully matched across the series; the consequences for the differential design are addressed in replicate-aware differential testing.

### snRNA-seq processing

#### Ambient RNA correction and quality control

CellBender remove-background was applied per sample on GPU with a false-positive rate of 0.01, 150 training epochs, a low-count threshold of 0, and per-dataset total_droplets and expected_cells. Per-sample quality control computes standard metrics with scanpy.pp.calculate_qc_metrics over mitochondrial (MT-/mt-prefix, species-aware), ribosomal, and hemoglobin gene sets, then filters cells on minimum detected genes, genes on minimum detecting cells, and cells on mitochondrial fraction. Doublets are detected with Scrublet run per sample; groups too small or too sparse for Scrublet to fit are retained unfiltered and logged rather than dropped. A full attrition ledger comprising filtered versus surviving cells, complete with justification is emitted per sample.

#### Integration

Samples are concatenated and integrated with scVI over the top 4,000 highly variable genes, using the manifest batch column as the batch covariate, for 50 epochs (PBMC, Brain, Kidney) or 100 epochs (AD).

#### Cell-type annotation

Three strategies are supported. By default, CellTypist assigns labels from a pre-trained reference model (Immune_All_Low.pkl for PBMC; Mouse_Whole_Brain.pkl for both mouse brain datasets). CellTypist models can also be trained from an adequately annotated reference, and the mouse kidney annotations demonstrate this process using the process outlined in the CellTypist documentation with our specific codes provided within our own repository documentation. Alternatively, marker-based annotation scores user-supplied marker sets with scanpy.tl.score_genes and assigns the top-scoring class subject to two guards: an absolute score floor (marker_min_score) and a minimum margin between the best and second-best score (marker_score_margin, default 0.1), below which the call collapses to *unknown* rather than resolving a near-tie arbitrarily. Finally, when a reference atlas directory is configured, scANVI is trained for label transfer over 50-100 epochs. As a demonstration for SCANVI functionality, the Allen mouse brain reference atlas v2 was used for annotation of the mouse AD dataset.

#### Differential expression

Per cell type, differential expression between condition groups is computed with MAST following conversion of the annotated AnnData object to a Seurat object. Positive log₂ fold-changes denote up-regulation in the treatment condition, matching the

<TREATMENT<_vs_

<CONTROL= output naming. These cell-level results are used for marker discovery and descriptive comparison; condition contrasts reported as inferential results are tested at the sample level, as described in replicate-aware differential testing.

#### Gene set enrichment

Gene Ontology and pathway enrichment is performed with enrichR against GO_Biological_Process_2023, GO_Cellular_Component_2023, GO_Molecular_Function_2023, and when applicable, KEGG_2021_Human; each is run separately on up-and down-regulated gene sets. For the AD condition contrast, enrichment is run twice. First on gene sets defined by the cell-level MAST contrast and then on gene sets defined by the sample-level edgeR contrast enabling direct comparison of the two units of analysis.

#### Co-expression networks

hdWGCNA is run per cell type, subject to the floors in Table S2. Genes are selected by expression fraction (gene_select = “fraction”, fraction = 0.05). Metacells are constructed with MetacellsByGroups grouped by cell type and sample, using the PCA reduction, *k* = 20, max_shared = 10, and a 50-cell floor, and are then normalized. Soft-thresholding power is chosen from a signed-network scale-free-topology sweep over powers 1-10 and 12-30 in steps of 2, taking the smallest power satisfying *R*^2^ > 0.8 and mean connectivity below 100, relaxing to mean *k* < 200 and then to maximum *R*^2^ if no power qualifies, with a hard floor of 3. Networks are constructed as signed, module eigengenes are computed with group.by.vars = “sample” where sample information exists, and intramodular connectivity is computed with ModuleConnectivity. Differential module eigengene testing and module-trait correlation constitute an optional second tier activated by setting the condition keys.

#### Cell-cell communication

CellChat is run on the annotated RNA object against the species-matched CellChatDB. Over-expressed genes and interactions are identified with do.fast = FALSE, communication probability is computed with the triMean method, and communications supported by fewer than 10 cells in either the sender or receiver population are filtered out before pathway-level aggregation. With cellchat.conditions populated, per-condition networks are inferred separately and contrasted; otherwise a single global network is produced.

### snATAC-seq processing

#### Two-pass quality control

ATAC_INITIAL imports fragments, computes per-sample fragment-count and transcription start site enrichment (TSSE) distributions using the dataset GTF, and emits data-driven thresholds. Initial filters are reported as tuples (initial min counts, initial min tsse), dataset-specific, yet intentionally permissive: PBMC (1000, 5), AD (1000, 2), Brain (500, 2), Kidney (500, 6). ATAC_FINAL applies the computed thresholds. The corresponding parameters params.atac.min_counts, min_tsse, and max_counts are null by default precisely so that they are derived rather than assumed; setting them explicitly overrides the computed values.

#### Per-sample processing

After threshold filtering, a 5-kb tile matrix is constructed with snap.pp.add_tile_matrix, the top 50,000 features are selected, and doublets are removed with SnapATAC2’s Scrublet implementation at a probability threshold of 0.5. A guard rejects the doublet call and retains all cells when every cell in a sample would be removed, assuming unreliable scores rather than a pure-doublet sample.

#### Dimensionality reduction and clustering

Samples are combined into an AnnDataSet, features are re-selected, and a spectral embedding is computed with snap.tl.spectral. Batch correction is configurable as none (used throughout this work), Harmony (max_iter_harmony = 20), or mutual-nearest-cell correction. A *k*-nearest-neighbor graph and UMAP are computed on the selected representation, and Leiden clustering is run at resolutions 0.5, 1.0, and 2.0, each stored under its own key.

#### Peak calling

Peaks are called per cluster with MACS3 through snap.tl.macs3 at the highest clustering resolution, filtered, and merged into a union peak set with snap.tl.merge_peaks against the genome chromosome sizes. A cell × peak matrix (snap.pp.make_peak_matrix) and a cell × gene activity matrix (snap.pp.make_gene_matrix) are then constructed.

#### Cell-type annotation

ATAC annotation operates on the peak matrix directly and is independent of the RNA arm. The default method is scATAnno reference-atlas annotation with a distance threshold of 95, an uncertainty threshold of 0.5, 30 dimensions, and 30 neighbors. A marker-based mode overrides this and writes to cell_type if desired. We note that imputed gene-activity scores may yield poorer annotations than scATAnno given assumed expression from chromatin accessibility is a more limited reflection of cell state.

#### Differential accessibility

Per cell type, differential accessibility between a treatment and a control condition is computed with snapatac2.tl.diff_test on the peak matrix, returning per-peak log₂ fold-change, p-value, and adjusted p-value. As for expression, these cell-level results are descriptive; condition contrasts reported as inferential results are tested at the sample level, as described in replicate-aware differential testing.

### Replicate-aware differential testing

Condition contrasts reported as inferential results were tested with the biological sample, not the cell, as the experimental unit. Cell-level tests are retained for descriptive one-versus-rest comparisons and marker discovery. Both modalities use a single framework, differing only in the count matrix drawn from and the design fitted.

#### Aggregation

Let *y*_g*i*_ be the raw count of feature *g* (gene for RNA, peak for ATAC) in cell *i*, and let *τ*(*i*) and *σ*(*i*) be its cell-type label and sample of origin. For each cell type *t* and sample *s*, pseudobulk profiles are formed by summing raw counts over the constituent cell set 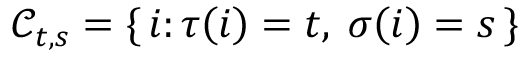:

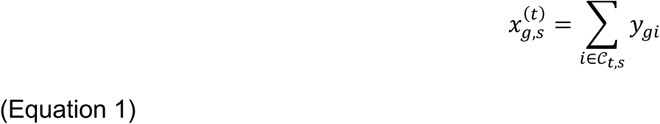

For RNA, the raw counts layer is summed. For ATAC, the fragment count matrix over the consensus peak set is summed. Columns are indexed by the pair (*t*, *s*), so within a cell type the analysis operates on a feature by sample matrix of at most twelve columns.

#### Inclusion

A pseudobulk column is retained only if it aggregates at least ten cells. A cell type is tested only if it satisfies the per-cell-type resolution floor and if at least three retained columns remain in each condition (Table 2). Cell types failing either criterion are excluded and logged.

#### Normalization and feature filtering

Composition bias between columns is corrected by the trimmed mean of M-values, giving effective library sizes *N*_j_ = *L*_j_*f*_j_ from observed library sizes *L*_j_ and normalization factors *f*_j_. Feature *g* is tested if CPM_gj_ = 10^6^*x*_gj_/*N*_j_ is at least 1 in no fewer than *k* = min(*n_TG_*, *n*_W*T*_) columns and its total count is at least 15.

#### Model

Filtered counts are modeled as negative binomial with a feature-specific dispersion *φ*_g_, a log link, design vector **z**_j_, and log*N*_j_ as an offset:

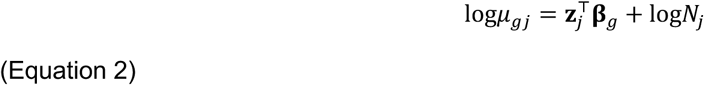

Dispersions are shrunk toward a robustly fitted abundance-dependent trend and a quasi-likelihood negative binomial generalized linear model is fitted, with the feature-level quasi-dispersion moderated by empirical Bayes toward that trend. The contrast is tested by the quasi-likelihood F-statistic

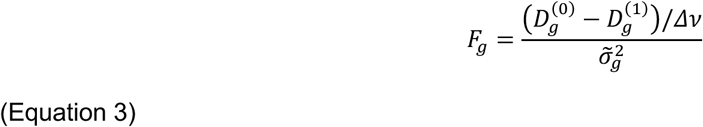

in which 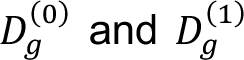 are the residual deviances of the null and full models, *Δν* the difference in their dimensions, and 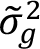 the moderated quasi-dispersion on *d* + *d* degrees of freedom, so that uncertainty in the dispersion estimate propagates into the test. The biological coefficient of variation is reported as *BCV_g_* = 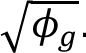.

#### Designs

Three contrasts are fitted, differing only in **z**_j_. For the RNA condition contrast, **z**_j_ carries an intercept, age in months as a continuous covariate, and a transgenic indicator as the tested coefficient; age is continuous rather than categorical because the oldest transgenic and wild-type ages are unmatched, and a factor encoding would restrict the condition estimate to the age-matched subset. For the ATAC condition contrast the age term is dropped. For the positive control, two transcriptionally distinct cell types measured in the same animals, **z**_j_ carries an intercept, one indicator per animal, and a cell-type indicator as the tested coefficient; because every animal contributes both cell types, the animal terms absorb between-animal variation and the contrast is within-animal. The reported effect size is log_2_FC_g_ = *β*_g,cond_/log2, positive values indicating higher abundance in the transgenic condition.

The age term was retained per modality on the basis of its effect on estimated dispersion, since a covariate explaining real variance reduces residual dispersion. Comparing the common BCV per cell type under both designs, the age term reduced it in every RNA stratum tested (mean reduction 18.0%) but in fewer than two-thirds of ATAC strata, increasing it in the remainder (mean reduction 5.6%).

#### Calibration and power

False-positive behavior was assessed by repeating the entire procedure under *B* = 50 balanced permutations of the condition label across the twelve animals, drawn without replacement from the 462 distinct six-versus-six splits and excluding the identity. Writing *S_b_* and *S*_0_ for the numbers of features declared significant under permuted and true labels, the empirical p-value is

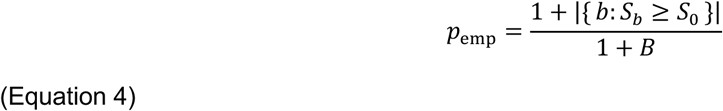

The same permutations were applied to a cell-level Wilcoxon rank-sum test, so that the two units of analysis could be compared on identical data. Sensitivity to sample size was assessed by subsampling animals without replacement. *n* = 2 to 6 per condition group for the condition contrast, and *n* = 2 to 12 in total for the paired control; with ten independent draws at each *n* except the largest, at which only one distinct subset exists. Curves are summarized by the median across draws with the full range shown, since the distribution of counts is strongly right-skewed at small *n*.

#### Correction

Within each cell type and modality, p-values are adjusted by the Benjamini-Hochberg procedure, the cell type being the unit at which each hypothesis family is defined. Features with an adjusted p-value at or below 0.05 are significant; no fold-change threshold is imposed.

Aggregation was performed in Python with h5py and NumPy, reading count matrices in row-blocked fashion to bound memory; testing was performed in R with edgeR, limma, statmod, and locfit. One aggregation task is emitted per modality x annotation and one test task per cell type.

### Multiome processing

#### Joint object construction

BUILD_MUDATA assembles the annotated RNA and ATAC objects into a joint MuData (.h5mu), processing samples in batches of mudata.batch_size (default 10) to bound memory. Barcode conventions differ between modalities and platforms (sample:barcode versus barcode-sample; 10x versus BD), so all cross-layer matching is performed in a normalized (sample, stripped barcode) space rather than on raw obs_names. Observation-column dtypes are sanitized before writing, and categorical columns with missing values are stored as pd.Categorical with a missing code of −1 rather than stringified, so that absent labels never become the literal string ’nan’.

#### MOFA+

Multi-omics factor analysis is run in high_memory mode with 15 factors and medium convergence. Factor stability is assessed by bootstrap, with consensus assessed by Spearman correlation of matched factors across bootstrap replicates.

#### MultiVI

A joint variational model is trained with scvi-tools for 200 epochs with a 20-dimensional latent space, equal modality weights, a Jeffreys modality penalty, and sample_id as the batch key. Because MULTIVI.train hard-codes an early-stopping patience of 50, a custom callback is supplied to expose and control early stopping. Three optional analyses are available and were run on selected datasets: a masking sweep that holds out 25%, 50%, and 75% of one modality and scores reconstruction of the held-out values; driver-factor analysis by Jacobian attribution, with Pearson correlation used for correspondence against MOFA+ factors; and gap-fill imputation for cells present in only one modality, using 25 posterior samples and a minimum confidence of 0.3. Reconstruction quality is scored as the per-gene Spearman rank correlation 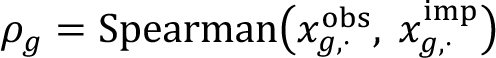 over the 3,000 highly variable genes, and reported as the distribution of *ρ*_g_ rather than a single pooled value, so that poorly reconstructed genes remain visible.

### Regulatory analysis

#### Co-accessibility

The Cicero stage is decomposed into distance-parameter estimation, per-chromosome model fitting, a join step, and peak-to-connection lookup against a chromosome-indexed dictionary. Cicero is run on the peak matrix through monocle3 with sample_num = 100 and num_dim = 50. Individual peak pairs are retained as arcs at a co-accessibility cut-off of 0.25, and connected components of the retained graph are assembled into co-accessibility modules (CCANs) subject to a minimum module co-accessibility of 0.1.

Three stratifications are available and are complementary rather than alternative. *Global* (the default) computes one co-accessibility map across the entire peak matrix; this map is neither per-cell-type nor condition-aware, and is reported as such. *Stratified* (the default for differential analyses) runs Cicero separately within each condition and contrasts the two maps with COMPARE_COACCESSIBILITY; it activates automatically whenever differential.run = true and a condition key is set. *Per cell type x condition* runs Cicero pairwise over (cell type, condition), subject to the 250-cell per-stratum floor in Table 2, which is a separate axis from the global per-cell-type floor. Strata below the floor exit with a dedicated code are skipped and logged.

#### TF motif enrichment and differential TF accessibility

GPU-accelerated chromVAR, implemented via scPrinter with a CuPy and RAPIDS RMM backend, computes per-cell motif deviation z-scores against JASPAR2022 motif sets, processing 30,000 peaks at a time. Per cell type, the top 5 motifs with a minimum absolute deviation z-score of 1.5 are retained; an optional global_top_n cap bounds the total number of unique transcription factors carried forward. These post-filtered factors are the primary gate on the scale of the footprinting that follows. Per-cell-type testing on the chromVAR deviation matrix runs in one of two modes: in *descriptive* mode each cell type is tested against all other cells, and in *differential* mode a treatment condition is tested against a control within a cell type. Both use scanpy.tl.rank_genes_groups with the Wilcoxon rank-sum test, use_raw = False, and reporting of expressing fractions, subject to the per-group floor in Table S2. Where a permutation contrast is required instead, scPrinter’s chromvar.permutation_test is used with 1,000 iterations.

#### TF footprinting

A scPrinter printer object is built from the ATAC fragments and cell-type barcode assignments. Multi-scale footprints and TF binding scores are computed against JASPAR2022 core non-redundant position frequency matrices at a 0.05 false-discovery-rate threshold. Promoter regions are defined as 2,000 bp upstream to 500 bp downstream of the transcription start site for footprinting; a separate 2,000/2,000 bp window is used when resolving coordinates for overlay motif scans, so that the scan covers the full gene-proximal regulatory region. Because printer objects are mapped to cell types, multi-scale footprinting is executed in a per-cell-type architecture in which the printer and peak matrix are loaded once per cell type rather than once per cell type-TF pair, reducing the overall task count, minimizing compute demands, and simplifying the fan-out.

#### Enhancer footprinting recipes

Candidate enhancers are defined in one of two ways: as members of a CCAN, or as peaks whose pairwise co-accessibility exceeds the 95th percentile of the observed distribution. The selected regions are scanned for TF motifs with MOODS via scPrinter against the same motif set, footprinted separately within each cell type, and quantified in two tables: one row per (cell type, TF) pair, and one row per (cell type, TF, gene) triple. The columns fall into four classes of evidence, ordered by the strength of the putative claim each supports: direct binding, cell-type specificity, enhancer-gene linkage, and differential activity. A provenance column, evidence_tier, records whether differential testing was possible for that cell type. Where a downstream step requires a binary rather than a continuous binding call, the threshold is set by Otsu’s method (Equation 10).

#### Domains of regulatory chromatin

Peak-gene correlations are computed with scPrinter’s fast_gene_peak_corr within a 250-kb window upstream and downstream of each gene. Significance is assessed against 500 background peaks matched per foreground peak, which controls for GC content and accessibility, yielding an observed correlation and a background-calibrated z-based p value; pairs with *p*_B_ ≤ 0.05 are retained. DORC genes are ranked by significant-peak count and the inflection point is visualized by J-plot; per-cell DORC scores are then computed to define the statistically significant set.

#### Topic modeling and eRegulons

The pycisTopic stage runs in three phases: per-group objects are constructed; a parallel latent Dirichlet allocation (LDA) sweep over topic counts of 10, 20, and 30 runs one job per count; and model selection plus binarization are executed. LDA treats the binary accessibility matrix as a document–term matrix and factorizes it into topic–region and cell–topic distributions, the former binarized in phase 3 to region sets and the latter used as a latent embedding, both sampled by MALLET’s collapsed Gibbs sampler over 500 iterations (23,52). FORGE passes --alpha 50.0 and --eta 0.1; because pycisTopic’s run_cgs_models_mallet defaults to alpha_by_topic=True and eta_by_topic=False, the effective priors are α_eff = 50.0/T and η_eff = 0.1, so the sweep varies the Dirichlet prior alongside the topic count (α_eff = 5.0, 2.5, and approximately 1.67 at T = 10, 20, and 30). Topic count is selected by evaluate_models(), which scores each candidate on topic coherence over its top 20 regions averaged across the top 5 topics (higher is better), topic redundancy as the mean pairwise cosine similarity between topics (lower is better), a symmetrized Kullback–Leibler divergence between the topic-region and cell-topic matrices (lower is better), and model log-likelihood (higher is better), each computed as defined in the originating publications (20,21); log-likelihood rises near-monotonically with T and is read alongside the other three rather than alone. FORGE forwards --selected-topics to select_model, and when that is unset the library’s choice stands.

Topics are binarized twice and both results retained: by Otsu’s method (method=“otsu”), which gives data-driven set sizes, and by fixed top-N (method=“ntop”, ntop=3000). For differential accessibility the accessibility matrix is reconstructed from the model as the product of the cell–topic and topic–region distributions and rescaled to a per-cell sum of 10⁴, with imputation restricted to the union of the binarized region sets rather than the full matrix. Region sets are tested against the cisTarget rankings and scores databases, which for mouse datasets on GRCm39 were regenerated from the aertslab mm10 v10nr_clust databases by UCSC liftOver, retaining 1,110,637 of 1,110,655 regions; recovery-curve area under the curve is normalized against the database-wide background to a normalized enrichment score and enriched motifs are mapped to transcription factors through the motif-annotation table (20,24,25,80), at ctx_rank_threshold 0.05, ctx_auc_threshold 0.005, ctx_nes_threshold 3.0, and orthologous_identity_threshold 0.0. SCENIC+ then ranks region-to-gene and TF-to-gene links by gradient-boosting feature importance, which is unsigned, taking direction from the Spearman correlation ρ between region accessibility and gene expression across cells with |ρ| < 0.05 discarded, and thresholds links in parallel by importance quantile (0.85, 0.90, 0.95), top-N regions per gene (5, 10, 15), and the binarized region sets. Direct eRegulons retain only regions carrying a directly annotated motif for the factor; extended eRegulons also admit motifs annotated by orthology or similarity and are scored separately into AUCell_extended.h5mu.

Regulon activity is scored per cell with AUCell over the top 5% of each cell’s feature ranking (auc_threshold = 0.05), separately for the gene-based and region-based signatures, so that each eRegulon carries both a transcriptional and an accessibility activity per cell; the score is rank-based and therefore invariant to library size and to monotone normalization. Cell-type specificity is summarized at both fine and broad annotation resolutions by the regulon specificity score, which compares an eRegulon’s AUCell profile p^_R across cells against the indicator vector q^_k for cell type k, both L₁-normalized:

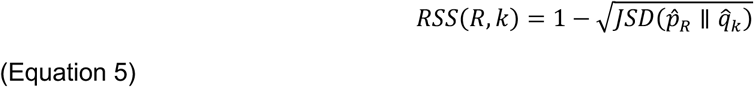

The square root makes this the Jensen-Shannon distance, as returned by scipy.spatial.distance.jensenshannon. Because scipy is called without base=2 the natural logarithm applies, bounding the distance by √(ln 2), so RSS ∈ [0.1674, 1], with 0.1674 rather than 0 indicating no specificity. RSS is computed by compute_rss in bin/plot_scenicplus_results.py, equivalent to scenicplus.RSS.regulon_specificity_scores except that regulons with an identically zero AUC vector return 0 rather than NaN.

#### ATAC-only TF-gene network layer

An optional layer (build_network) combines continuous scPrinter binding scores with Cicero co-accessibility to link transcription factors to target genes without requiring RNA. This is recommended for gene regulatory network construction only when running in standalone ATAC mode, given the robustness that RNA information grants to DORC and SCENIC+.

### Visualization

FORGE renders figures as pipeline outputs. Coverage tracks are exported per pairwise cell type-condition at 10-bp bins, subject to the floor in Table S2. Genome-browser panels are rendered matplotlib-natively with a default 15-kb padding at 700 bins, auto-scaling the y-axis to the 99.5th percentile by default; this affects display only, not the underlying values. Cicero target-gene arc plots are rendered at three mirrored upstream and downstream threshold tiers simultaneously: 100 kb at 0.10, 250 kb at 0.05, and 500 kb at 0.025.

Composite enhancer panels are restricted by an explicit candidate filter for genes, transcription factors, and cell type. Multi-scale footprint strip plots for promoters and enhancers render in absolute, differential, or combined mode. Enhancer-strip target genes are discovered rather than configured, ranked per pairwise cell type-TF from TF binding scores crossed with Cicero co-accessibility over the top 100 enhancer peaks, retaining the top 5 genes.

An additional figure suite (SHI_FIGURES) mirrors the analysis panels of our previous work (103): peak-type annotation against a 2-kb promoter window, non-negative matrix factorization enhancer programs over a *k*-grid subject to the floor in Table S2, marker coverage tracks, co-accessibility correlation matrices, differential-accessibility breakdowns and log₂ fold-change heatmaps, TF differential volcano plots, curated TF networks, and locus-level TF binding panels. Tier B panels are gated on condition labels and depend on upstream differential accessibility results.

### Three-tier verification strategy

#### Tier 1: structural verification

Tier 1 verifies the Nextflow architecture and is executed with nextflow run main.nf -profile test -preview -c configs/datasets/test_preview.config against a self-contained test_data/fixture included in the repository. The test profile strips every site-specific scheduler assumption, disables containers, and redirects output to results_test/. The -preview flag constructs the complete process graph and executes the full pre-flight checklist without submitting any task, reporting a pass after nine checks.

#### Tier 2: small real run

A CPU-only tutorial configuration executes real tools on a subsampled 10x PBMC dataset with ATAC fragments restricted to chr21 and chr22, taking 20,000 input barcodes into RNA processing, consisting of 1,000 cells and 19,000 background barcodes. chromVAR is excluded from this tier because bin/gpu_chromvar_nf.py imports CuPy and RAPIDS RMM at module scope; SCENIC+, pycisTopic, and scPrinter footprinting are also excluded, because cisTarget databases cannot be cheaply subset and footprinting accounts for the majority of pipeline compute. All three are instead demonstrated through the on-ramp mechanism with pre-computed outputs. Tier 2 is not intended to produce results appropriate for biological interpretation; it is intended to demonstrate quickly that genuine biological data moves predictably through the FORGE architecture. Cell, gene, and peak counts surviving each stage, and total runtime, are reported in results.

#### Tier 3: full run

Tier 3 is FORGE with full capabilities enabled and describes the state of the four validation datasets in Table S3.

### Compute accounting

Pipeline cost was measured across all four datasets by joining three independent provenance sources per task: .nextflow.log* (task runtime, complete across resume sessions), trace.tsv (the Nextflow cpus directive, requested memory and peak resident set size (RSS); a single-session snapshot) and SLURM sacct (time limit, requested and allocated CPUs, requested memory, elapsed time, maximum RSS, total CPU time and billing TRES). Log and trace records were joined on the Nextflow work-directory hash and to sacct on the SLURM job ID, the hash also serving as the deduplication key so that each row is one distinct task execution; because the union spans resume sessions, retried tasks appear as separate executions and the totals are an upper bound on a single clean run. The ledger comprises 4,984 unique executions, all resolving to COMPLETED or CACHED in the retained logs. Over tasks *i* with realtime *tı* (hours):

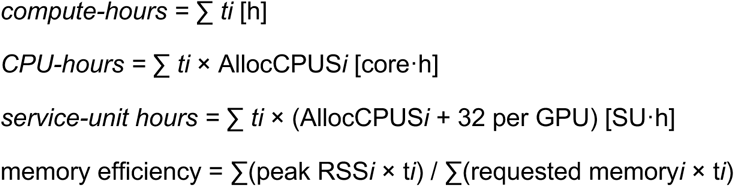

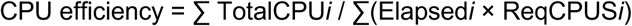

CPU-hours use allocated rather than directive CPUs because this cluster satisfies a memory request by allocating additional cores, and both memory-efficiency terms are weighted by realtime so that deliberate walltime over-request is not reported as a memory result; peak RAM, a maximum rather than an additive quantity, is reported per process as the maximum over tasks of max(trace peak RSS, sacct MaxRSS). Given the aggregate is dominated by an opt-in layer, cost is reported in three cumulative tiers comprising the core pipeline, advanced regulatory, and full FORGE. These designations were assigned structurally from the fully-qualified Nextflow process name together with the pipeline’s own gating parameters (Figure 2A), and every per-cell-type step is additionally re-costed as mean per-task runtime × number of cell types to estimate scaling over one clean pass (Figure 2C).

### Post-hoc evaluation analyses

Post-hoc analyses were performed outside the pipeline, operating on persisted pipeline outputs, in order to facilitate evaluation across the datasets. More detailed assessments against external benchmarks were performed for the public human PBMC and mouse TgCRND8 datasets, where literature was available for comparison. Throughout, *n* denotes the number of cells, *c* indexes cell-type classes, and *k* = 15 is the neighbourhood size for every *k-*nearest-neighbor quantity defined in this section (layers L2 and L3). These scores build their own exact neighbor graphs from the persisted embeddings, excluding each cell itself, and are therefore independent of the neighbor graphs constructed inside the pipeline, which use separately chosen sizes. Explicitly, 30 for scANVI-based RNA clustering and MultiVI UMAPs, 15 for MOFA UMAPs, and the SnapATAC2 default of 50 for ATAC clustering.

#### Cross-modal concordance

Because FORGE annotates RNA and ATAC independently, their agreement is measurable, and concordance was quantified in three layers on all four datasets.

Layer L1 assesses raw RNA-ATAC cell-type agreement from the per-cell confusion matrix over broad labels. The matrix *N* has entries *N_ab_* counting cells labeled *a* in RNA and *b* in ATAC; cells on which the two annotations agree fall on the diagonal, and L1 is the fraction of jointly classifiable cells lying there:

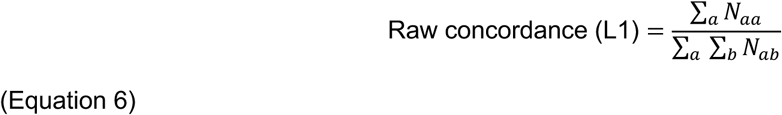

Layer L2 evaluates embedding cohesion, separately under RNA and ATAC labels, in four latent spaces: the MultiVI latent dimensions, the MOFA+ factor space, raw RNA principal components, and the raw ATAC spectral embedding. Cohesion measures whether cells sharing a label are neighbors in a latent space. For each cell *i*, let *N_k_*(*i*) be the set of its *k* nearest neighbors in that space, excluding *i* itself, and let *l*(*i*) denote its label; writing *j* for a neighbor in that set, cohesion is the mean fraction of neighbors sharing the cell’s own label:

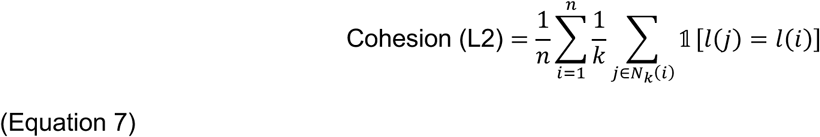

Because cohesion has a non-zero chance level that differs between datasets, raw values are accompanied by a scaled score, (observed − floor)/(ceiling − floor). The chance floor is the *k*-nearest-neighbor purity null computed from the dataset’s own label marginal, floor = 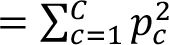 with *p_c_* = *n_c_*/*n, C* the number of broad classes, and *n_c_* the number of cells in class *c*. Two floors are computed per dataset from the L1 confusion matrix: the RNA floor from its row marginal, applied to the RNA cohesion block, and the ATAC floor from its column marginal, applied to the ATAC cohesion block. The home-court ceiling is the modality’s cohesion in its own raw space. That is RNA cohesion in raw RNA principal components for the RNA block, and ATAC cohesion in the raw ATAC spectral embedding for the ATAC block. The resulting scale reads 0 at chance and 1 at the modality’s own home space. This is the scIB and GLUE scaled-score paradigm; scaling against the ceiling alone would launder near-chance results into high fractions. Values above 1.0 are reported as the joint embedding sharpening structure and are not clipped.

Layer L3 evaluates cross-modal label transfer accuracy in each of the L2 spaces. Within each space, each cell’s label is predicted by majority vote over its *k* nearest neighbors drawn from the other modality’s labeled cells, and accuracy is the fraction of cells whose transferred label matches their own:

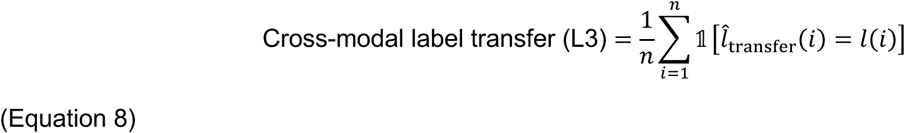

L3 is read against the L1 row rather than rescaled, since transfer accuracy is bounded by the raw agreement it attempts to reproduce.

#### External comparison for the PBMC dataset

Three comparison axes were constructed for the human PBMC dataset, the only one of the four for which external quantitative benchmarks exist.

##### Axis 1: annotation concordance against a reference-mapped standard

No published cell-type labels exist for this 10x PBMC multiome dataset, so an external standard was generated rather than adopted. The previously published multimodal PBMC reference was loaded from its deposited object and the FORGE-processed PBMC RNA object was projected into it with Seurat FindTransferAnchors and TransferData, transferring reference level-2 labels onto the query cells (104). Transferred labels were cross-tabulated against FORGE’s broad and fine annotations and summarized by adjusted Rand index, adjusted mutual information, and homogeneity. Adjusted Rand index is computed over all 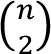 cell pairs, with *N_i_*_j_ the contingency table between the two labelings and *a_i_* and *b*_j_ its row and column margins:

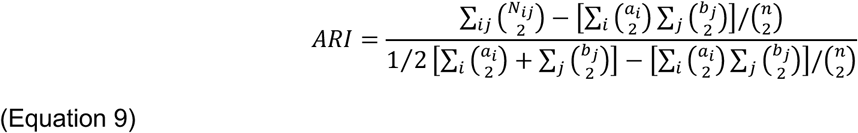

ARI is 0 at chance agreement and 1 at identity, and may be negative for agreement worse than chance. For adjusted mutual information, with *U* and *V* the two labelings, entropy *H*(*U*) = − ∑*_i_ P* (*i*)log*P*(*i*) with *P*(*i*) = |*U_i_*|/*n* measures the uncertainty in a random cell’s label, and mutual information

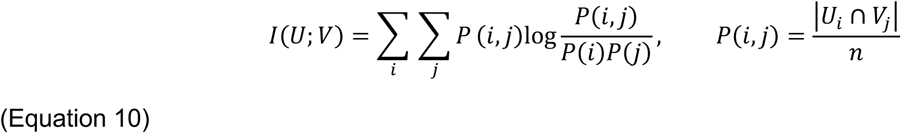

measures how much knowing a cell’s class in one labeling determines its class in the other; it is 0 when the labelings are independent and equal to *H*(*U*) when they are identical. Because mutual information rises with cluster count irrespective of agreement, it is adjusted against the value expected from random labelings with the same class sizes, computed under the hypergeometric model:

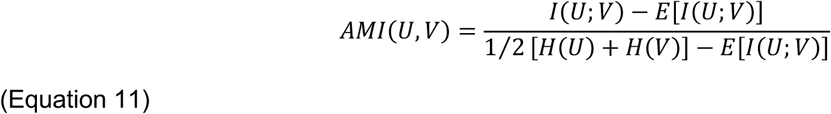

The arithmetic mean of the two entropies is used as the normalizer, and AMI is 0 at chance agreement and 1 at identity. Homogeneity, ℎ = 1 − *H*(*U* ∣ *V*)/*H*(*U*) with *U* the transferred reference labels and *V* the FORGE labels, measures the extent to which each FORGE cluster contains cells of a single transferred-label class; it is 1 when every FORGE cluster is pure with respect to the reference labels and 0 when the labeling is uninformative. Unlike ARI and AMI, which are symmetric in their two arguments, homogeneity is asymmetric, and is reported alongside them. The per-cell mapping-score distribution is reported alongside all three so that low-confidence transfers remain visible.

##### Axis 2: cross-modal integration error against a published baseline

Fraction of samples closer than the true match (FOSCTTM) was computed on FORGE’s MultiVI embedding by encoding each cell’s RNA and ATAC profiles separately and measuring, per cell and per direction, the fraction of all other cells lying closer than that cell’s own true match under a Euclidean distance *d*:

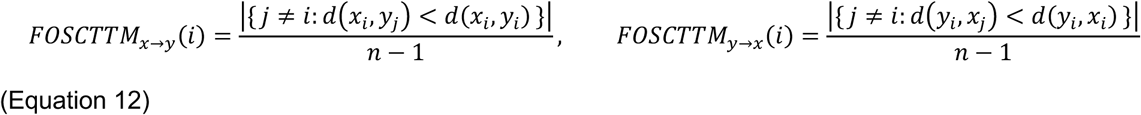

The reported score is the mean over all cells and both directions. Two identical embeddings approach 0, and two independent random embeddings approach 0.5, since the true match then holds a random rank. The published GLUE eleven-method baseline for this dataset was taken from that paper’s supplementary data; GLUE was not re-run. FOSCTTM was selected as the shared evaluation metric because published calculations exist for GLUE and because it is invariant to the composition of the benchmarked method set, whereas composite integration scores reported alongside it are min-max pooled over that set and change when a method is added or removed. FOSCTTM recomputed on a subsample matched to the GLUE cell count, and on artificially unpaired models from the masking sweep, is reported in results. The interpretive limits of this comparison are addressed in limitations of the study.

*Axis 3: regulatory corroboration.* Two PBMC-relevant regulatory findings reported by GLUE were interrogated against three independent FORGE layers: SCENIC+ eRegulons, DORC peak-gene links, and TF motif enrichment within significant DORC peaks tested against that motif’s genome-wide base rate. The base rate is taken as the fraction of all regions in the factor’s cistrome, so that a motif frequent across the genome is not counted as enriched merely for being common. Effect size is reported as enrichment = log_2_(*f*_peaks_/*f*_genome_), where *f*_peaks_ is the fraction of tested DORC peaks hosting the motif and *f*_genome_ the base rate. Significance is assessed by a one-sided exact binomial test of *n*_hit_ successes in *n*_peaks_ trials against success probability *f*_genome_, under the alternative that the peak set is enriched.

#### Cis-rewiring in the AD dataset

The glial *Spi1*/*Mef2c* disease program in the AD model was decomposed into a candidate *cis*-regulatory element rewiring component and a transcriptional component, evaluated per cell type and per condition. Co-accessibility arcs and CCANs are as defined in regulatory analysis. Differential accessibility and differential expression are both taken from the sample-level edgeR framework described above, so that the differential promoter-peak ATAC and the differential expression of the corresponding gene are compared one-to-one at the same unit of analysis, by Spearman rank correlation. The two coefficients come from differently specified models. The RNA design includes the age covariate and the ATAC design does not. This is further justified in that section and the correlation is interpreted accordingly.

Motif scanning used MOODS via scPrinter over the full 332,614-peak ATAC set with JASPAR2022 core non-redundant mm10 motifs, so that motif coordinates fully cover any CCAN enhancer. A position weight matrix is scored against a uniform background model, and a hit is called where the log-odds score exceeds the scanning threshold:

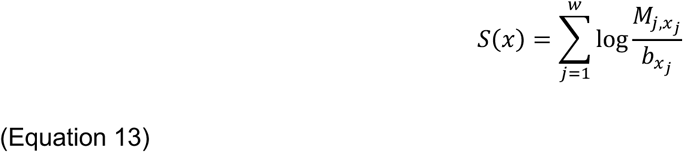

with *M* the position weight matrix, *w* the motif width, *x*_j_ the base at position *j*, and *b_x_*j = 1/4 for all bases under the uniform background.

##### Size-robust readouts

Cicero produces denser co-accessibility from smaller homogeneous populations. The reported inferences rest on four quantities that are relative or normalized and therefore robust to this confound; absolute network-size comparisons are treated as descriptive only.

First, motif fraction: of the enhancers gained in transgenic relative to wild-type animals, the fraction hosting a given TF motif, *f*_TF_ = *n*_gained enhancers with motif_/*n*_gained enhancers_, compared against the lineage background fraction as log_2_(*f*_TF_/*f*_bg_). Numerator and denominator both scale with network size, so the ratio cancels the cell-count confound.

Second, anchor-category proportion: each retained arc is classified by the peak-type pair of its two anchor peaks (promoter, exonic, intronic, distal) and reported as a proportion of all arcs within a condition, *P*_cat_ = *n*_arcs in category_/*n*_arcs total_. Comparing transgenic and wild-type proportions rather than counts cancels both cell-count and sequencing-depth effects.

Third, gene-set convergence: because more than half of enhancers are gained in transgenic animals, a binary gained-or-not classification is non-selective. Genes are instead ranked by the difference in link count between conditions, and the top-ranked genes tested for enrichment against three independent gene sets — the MEF2C SCENIC+ eRegulon targets, the SPI1 eRegulon targets as microglial ground truth, and DORC genes — by hypergeometric test against the linked-gene universe. Let *N* be the linked-gene universe, *K* the number of universe genes belonging to the target set, *N*_top_ the number of top-ranked genes taken, and *x* the number of target-set genes observed among them. The probability of drawing at least *x* target genes by chance, sampling without replacement, is

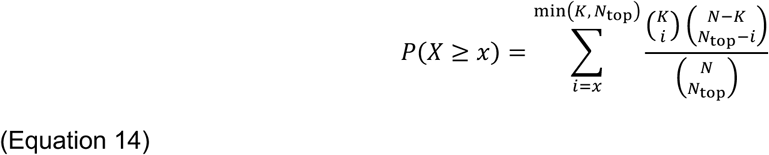

Each term is the probability of exactly *i* target genes among the *N*_top_ drawn. A small value indicates that the rewired genes overlap the target set more than chance allows.

Fourth, cross-type specificity: the same statistic computed across cell types, establishing that a signal attributed to one glial population is not present at comparable magnitude in the others. Specificity is read as the contrast between the focal cell type’s enrichment and the distribution across the remaining types, separating a cell-type-restricted program from a global one.

##### Positive controls

Two positive controls were run so that a null result is informative. First, lineage-marker cis-network specificity confirms that marker loci recover their expected cis-networks. Second, a cell-type contrast was run through the same pseudobulk edgeR quasi-likelihood machinery, filter, and twelve animals as the genotype contrast, differing only in its design. Given every animal contributes both cell types, the contrast blocks on animal (∼ animal + cell type) and is therefore within-animal, whereas the genotype contrast is necessarily between-animal. On the accessibility arm, immune versus OPC/oligodendrocyte returned 33,759 of 65,079 peaks at FDR < 0.05 (51.9 per cent; 29,350 at |logFC| > 1; BCV 0.133); on the expression arm, microglia/PVM versus oligodendrocytes returned 8,926 of 13,347 genes (66.9 per cent). Against these, TG versus WT returned 76 of 943,015 peak-level and 340 of 219,519 gene-level tests. Both modalities detect large effects at n = 6 per group; the genotype contrast is not one.

## QUANTIFICATION AND STATISTICAL ANALYSIS

### Software and multiple-testing correction

All statistical analysis was performed inside the five versioned Singularity images listed in Table S1; package versions are given in the key resources table. Multiple-hypothesis correction is Benjamini-Hochberg except where a package returns its own adjustment, which is noted per test below; it is applied through statsmodels.stats.multitest.multipletests(method=’fdr_bh’), scanpy’s built-in adjustment in rank_genes_groups, or the native adjustment of the R package in question. Unless stated otherwise, significance is defined as an adjusted p value below 0.05.

### Units of analysis

Three distinct units of *n* appear in this work.

- **Cells (nuclei)** — the unit for per-cell tests: cell-level differential expression and accessibility, differential TF accessibility, *k*-nearest-neighbor concordance metrics, and FOSCTTM.
- **Samples (animals or donors)** — the unit for pseudobulk analyses. The AD case-control pseudobulk analyses used *n* = 6 transgenic and *n* = 6 wild-type samples per cell type, subject to the inclusion thresholds in Table S2.
- **Features** — peaks, peak-gene pairs, genes, motifs, or (cell type, TF, gene) triples, as the unit for enrichment and correlation analyses.

Every reported condition contrast states which unit supports it. Where both units were computed for the same contrast, both are reported and the sample-level result is the inferential one.

### Randomization, blinding, sample-size determination, inclusion and exclusion

This is a computational methods study on existing and public datasets. No experimental randomization, allocation concealment, or blinding was applicable, and no animals or subjects were assigned to interventions by the authors. Exclusions are applied at defined, logged stages, and every excluded entity is recorded as attrition in a log file. The AD dataset’s transgenic and wild-type groups are the genotype groups of the source cohort.

Sample sizes were not determined by a prospective power calculation; the four validation datasets were selected to span the axes over which the pipeline was required to generalize. Namely, species (human, mouse), assay chemistry (10x Chromium, BD Rhapsody), sample number (1 versus 12), and design (single-condition versus case-control). Statistical power for the one case-control analysis is addressed directly by the positive control described in method details rather than by an assumed effect size: the same sample-level machinery applied to a cell-type contrast recovers abundant significant peaks where the genotype contrast recovers very few, which argues against a general analytical failure while not excluding limited power for subtler genotype effects. Counts for both contrasts are reported in results.

Seeds are as given in reproducibility, provenance, and random seeds.

### Statistical tests by analysis

#### Differential gene expression, cell level

MAST via Seurat::FindMarkers(test.use = “MAST”), applied per cell type, with cells as the unit. Reported per gene: average log₂ fold-change (positive denotes up-regulation in the treatment condition), unadjusted p value, and Bonferroni-adjusted p value as returned by Seurat — this is the one test whose correction is not Benjamini-Hochberg. Significance threshold: adjusted p < 0.05, with a log₂ fold-change cut-off of 0.25 applied for downstream gene-set selection.

#### Differential expression and accessibility, sample level

Negative binomial quasi-likelihood F-test in edgeR on pseudobulk profiles, per cell type, with the sample as the unit; designs, thresholds, and calibration as given in replicate-aware differential testing. Significance: adjusted p < 0.05, with no fold-change threshold. Reported per contrast: the number of significant features of the number tested, and the log₂ fold-change distribution.

#### Differential chromatin accessibility, cell level

snapatac2.tl.diff_test per cell type, treatment versus control, on the union peak matrix. The unit is the peak, and the number of peaks tested is reported per contrast. Significance: adjusted p < 0.05.

#### Cluster and cell-type marker genes

scanpy.tl.rank_genes_groups with the Wilcoxon rank-sum test, one cluster or cell type against all remaining cells.

#### Differential TF motif accessibility

scanpy.tl.rank_genes_groups with the Wilcoxon rank-sum test on the chromVAR deviation matrix, in descriptive or differential mode as described in method details; the unit is the cell. Where a permutation contrast was used instead, p values are corrected across motifs with multipletests(method=’fdr_bh’).

#### Gene set and motif enrichment

Enrichr p values are Fisher exact with Benjamini-Hochberg correction as returned by the service. For TF-motif over-representation in a foreground peak set relative to a background set, a one-sided Fisher exact test (scipy.stats.fisher_exact, alternative=’greater’); where the comparison is against a genome-wide base rate, a one-sided binomial test; and for gene-set convergence between rewired-gene rankings and eRegulon or DORC membership, a hypergeometric test (Equation 20). The unit is the feature.

#### Peak-gene association

DORC peak-gene correlations via scPrinter fast_gene_peak_corr; the unit is the peak-gene pair, and significance is background-calibrated as described in method details.

#### Correlations

Spearman rank correlation (scipy.stats.spearmanr) is used throughout: footprint against TF expression, footprint against target-gene expression, and TF against target in cross-modal validation and signal-chain analysis; observed against imputed expression per gene in the MultiVI masking sweep; promoter differential ATAC against differential expression in the cis-rewiring analysis; and factor matching across MOFA+ bootstrap replicates. Pearson correlation (scipy.stats.pearsonr) is used for MultiVI against MOFA+ latent-factor correspondence, for MultiVI imputation against observation, and for control against treatment co-accessibility scores over shared peak pairs, the last reported with its p value and the number of shared pairs. For the concordance between cell-level (MAST) and replicate-aware (edgeR quasi-likelihood) log₂(fold-changes), correlation is reported as both Pearson r and Spearman ρ, pooled and per cell type.

#### Annotation concordance and embedding metrics

ARI, AMI, homogeneity (Equations 9-11), cohesion and its scaled score (Equation 7), cross-modal label transfer (Equation 8), and FOSCTTM (Equation 12), all as defined in post-hoc evaluation analyses. Concordance statistics are reported with the full contingency table in both raw-count and row-normalized form. FOSCTTM is cell-count dependent, since more cells means more competitors, so the *n* used is stated wherever a value is quoted.

#### Co-expression networks

hdWGCNA soft-thresholding power selection as described in method details. Differential module eigengene testing uses the Wilcoxon rank-sum test (test.use = ’wilcox’) with the package’s adjusted p value and significance at adjusted p < 0.05; the unit is the cell, and the consequences are addressed in limitations of the study.

#### Enhancer program decomposition

Non-negative matrix factorization (sklearn.decomposition.NMF) on pseudobulk enhancer accessibility over a component grid of *k* = 6, 8, 10, and 12, with *k* selected by reconstruction-error elbow unless fixed.

#### Cell-cell communication

CellChat communication probabilities computed by the triMean method with the interaction filter given in Table S2. Where receiver-cell signaling scores were contrasted against other cell types, a one-sided Mann-Whitney U test (alternative=’greater’).

### Descriptive statistics and dispersion

Distributions are summarized by median and interquartile range where skewed (quality-control metrics, footprint statistics, and per-cell mapping scores) and by mean where a symmetric summary is appropriate (bind_dip_depth_mean, fp_depth_mean), with the 95th percentile reported alongside the mean for footprint dip depth. Violin and box plots show the median and interquartile range, with whiskers as specified in each legend. Correlation coefficients are reported with their p-value and *n*. Reference-mapping confidence is reported as the distribution of per-cell mapping scores, giving mean, median, and the fraction below 0.5.

### Assumption checking and robustness

The Non-parametric rank-based Wilcoxon rank-sum, Mann-Whitney U, and Spearman tests were used as the default for per-cell comparisons of expression, accessibility, and deviation scores, because these distributions are zero-inflated, non-normal, and no distributional assumption is warranted. Where a parametric model is used, it is one whose assumptions are explicitly modeled: MAST models the bimodal, zero-inflated expression distribution, and the edgeR quasi-likelihood framework models the mean-variance relationship of pseudobulk counts. The calibration of the sample-level test was assessed empirically by label permutation rather than assumed, and its sensitivity to sample size by subsampling, both as described in method details. Peak-gene correlation significance is not assessed against an analytic null but against 500 matched background peaks per foreground peak, which controls for GC content and accessibility.

Finally, all comparisons against published results are framed as cross-method concordance or corroboration. No external reference used in this work constitutes ground truth, and none is presented or intended to be interpreted as such; neither label vector in any concordance analysis is treated as truth, and no axis is labeled as such.

## ADDITIONAL RESOURCES

FORGE documentation and source code. Full user documentation, including a quickstart, chapters on the three files a user actually edits (the manifest CSV, the dataset config, and the main.nf architecture), installation and container-build instructions, reference-data preparation, per-arm how-to guides (RNA, ATAC, regulatory analysis, multiome integration, visualization), on-ramp and resumption recipes, cluster-adaptation guidance, and troubleshooting. The complete Nextflow pipeline, analysis scripts, container definition files, resource-tier configurations, and dataset configuration templates, released under a BSD 3-Clause license: https://swaruplabUCI.github.io/FORGE/

## Notes

https://github.com/swaruplabUCI/FORGE

https://swaruplabuci.github.io/FORGE/

### Competing Interest Statement

The authors have declared no competing interest.

